# Bacteriophage genome-wide transposon mutagenesis

**DOI:** 10.1101/2025.11.23.690004

**Authors:** Alex Chan, Wearn-Xin Yee, Deepto Mozumdar, Claire Kokontis, Matias Rojas-Montero, Li Yuping, Ying Yang, Joseph Bondy-Denomy

## Abstract

Bacteriophages have genomes that span a wide size range, are densely packed with coding sequences, and frequently encode genes of unknown function. Classical forward genetics has defined essential genes for phage replication in a few model systems but remains laborious and non-scalable. Unbiased functional genomics approaches are therefore needed for phages, particularly for large lytic phages. Here, we develop a phage transposon sequencing (TnSeq) platform that uses the *mariner* transposase to insert an anti-CRISPR selectable marker into phage genomes. CRISPR-Cas13a–based enrichment of transposed phages followed by pooled sequencing identifies both fitness-conferring and dispensable genes. Using the *Pseudomonas aeruginosa*-infecting nucleus-forming jumbo phage ΦKZ (280,334 bp; 371 predicted genes) as a model, we show that ∼110 genes are fitness-conferring via phage TnSeq. These include conserved essential genes involved in phage nucleus formation, protein trafficking, transcription, DNA replication, and virion assembly. We also isolate hundreds of individual phages with insertions in non-essential genes and reveal conditionally essential genes that are specifically required in clinical isolates, at environmental temperature, or in the presence of a defensive nuclease. Phage TnSeq is a facile, scalable technology that can define essential phage genes and generate knockouts in all non-essential genes in a single experiment, enabling conditional genetic screens in phages and providing a broadly applicable resource for phage functional genomics.

## Introduction

Bacteriophage genomes are rich in genes of unknown function and exhibit extensive genetic and proteomic diversity. Many fundamental discoveries likely remain hidden in uncharacterized phage genes^1^. However, few tools exist to enable unbiased genome-wide studies to bring phages into the functional genomics era. Most challenging perhaps are large lytic phages (e.g. jumbo phages with genomes >200 kb), where most genes are of unknown function^2^. For most of these phages, even the basic inventory of genes essential for replication is unknown.

Here we focus on ΦKZ, a model member of the *Chimalliviridae* family with distinctive infection biology^3,4^. ΦKZ has many notable features, including two distinct multi-subunit RNA polymerase complexes^5–8^, a lipid-based early phage infection vesicle^9–11^, and a protein-based phage “nucleus”^12,13^ that enables resistance to nucleases^14^. It remains unclear which genes are strictly required for this core biology, and which are dispensable but conditionally important, for example in specific hosts or environments. A method that could both rapidly identify all essential genes and simultaneously generate a library of knockouts in all other genes would greatly accelerate mechanistic dissection of such phages.

Targeted phage genome manipulation tools such as homology-directed recombination (HDR) and CRISPR-Cas counter-selection have been widely used^15^ albeit at low throughput. Recombineering is also effective and has recently been adapted to genome-wide mutagenesis^16^ and *in vitro* genome manipulation and rebooting have expanded the toolkit further^17^. RNA-targeting Cas13 has been particularly useful for selecting for spontaneous^14,18^ or HDR-derived phage mutants^19^. To enable facile and scalable bacteriophage genome modification with the ease afforded to bacteria by transposon insertion sequencing (TnSeq), we sought to develop an analogous genome-wide insertional mutagenesis method for phages. Here, we deploy *mariner* transposition in the nucleus-forming jumbo phage ΦKZ, using an anti-CRISPR (*acrVIA1*) selectable marker and CRISPR-Cas13a–mediated enrichment to isolate thousands of independently transposed phages. High-throughput mapping of insertion sites reveals essential genes at genome scale and yields a large collection of phage mutants for conditional phenotypic screens.

Genome-wide knockout approaches that generate permanent changes in phage genomes, such as the one presented here, are complementary to recently developed protein knockdown approaches based on antisense oligomers (ASOs) and dCas13d-mediated translation interference (CRISPRi-ART)^20,21^. These perturbation tools are powerful for studying essential genes that cannot be deleted outright. By contrast, phage TnSeq requires no large-scale oligo or guide RNA design, *mariner* transposase is active in many bacterial hosts^22^, and inserts transposons semi-randomly^23,24^. Together, these features should enable broad adoption of phage TnSeq as an easy-to-implement, low-cost genome-wide mutagenesis platform which can, in principle, be executed with any combination of a strong phage defense system and cognate anti-defense molecule.

## Results

### Establishing transposition in a nucleus-forming jumbo phage

ΦKZ protects its genome within a protein-based nucleus-like structure during infection and selectively imports proteins. To enable transposition inside this compartment, we fused the Mariner transposase to an sfCherry fluorescent reporter and to the nucleus-localized phage protein Nlp1^12,25^ (gp152) (Figure 1A). Upon induction of this fusion and infection with ΦKZ, fluorescence microscopy revealed co-localization of sfCherry–Mariner–Nlp1 with DAPI-stained phage DNA, indicating import of the transposase into the phage nucleus (Figure 1B).

**Figure 1.**
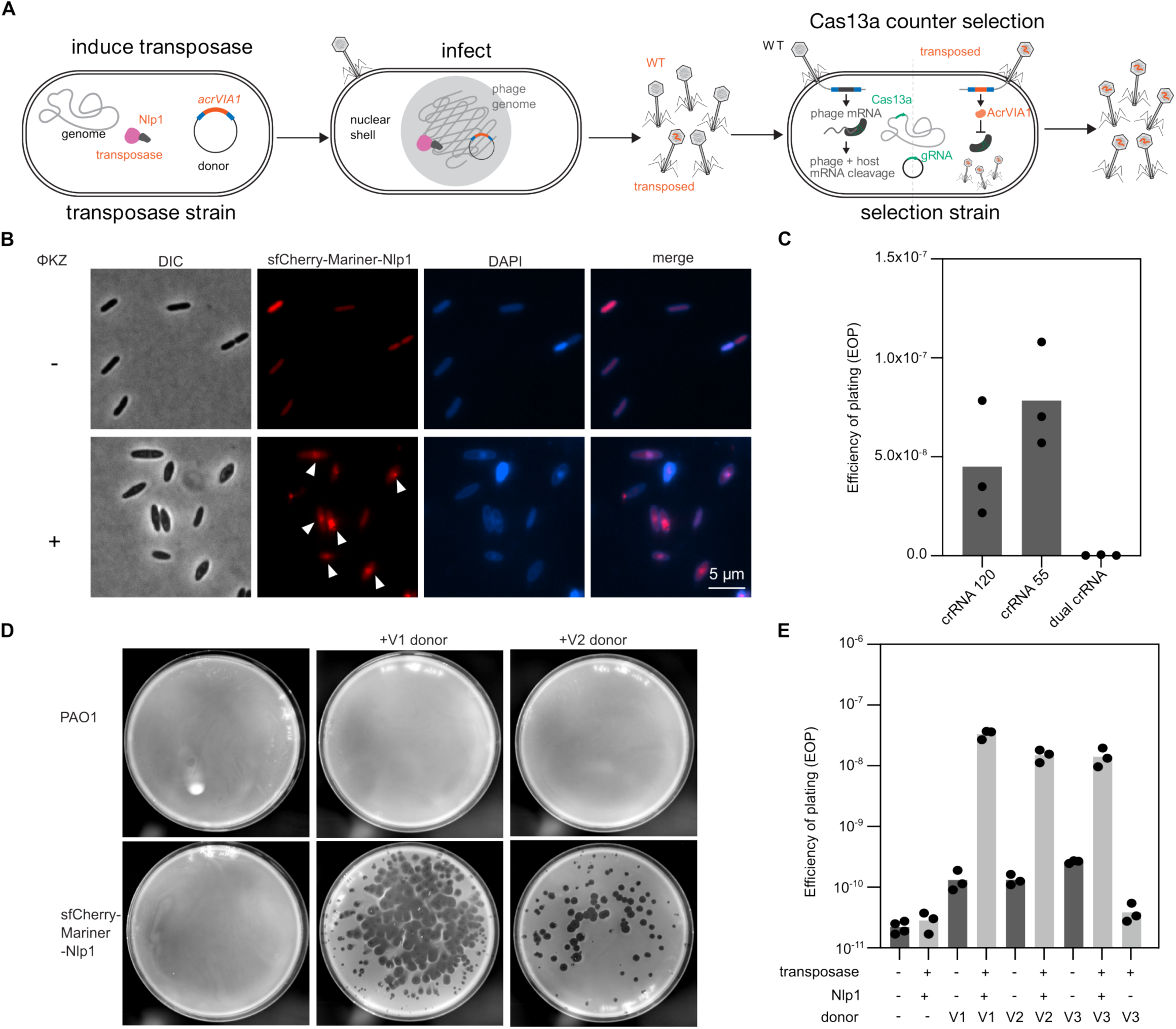
Transposon mutagenesis of a nucleus-forming jumbo phage. (A) Schematic illustrating the ΦKZ transposon mutagenesis strategy. The Nlp1–Mariner transposase fusion localizes to the phage nucleus, where a transposon encoding *acrVIA1* inserts into the phage genome, generating insertional mutants. Output phages are subsequently subjected to Cas13-based counter-selection. (B) Fluorescence microscopy of *P. aeruginosa* infected with ΦKZ expressing sfCherry-tagged transposase (red) and stained with DAPI (blue). White arrows indicate co-localization of transposase with the phage nucleus in infected cells. (C) Quantification of ΦKZ replication efficiency in the presence of single versus dual Cas13a crRNAs. (D) Representative plaque phenotypes of transposed ΦKZ forming on dual Cas13a targeting strains. (E) Efficiency of plaquing (EOP) of ΦKZ on dual Cas13a strains following transposition with variants V1 (*acrVIA1* with a phage early promoter), V2 (*acrVIA1* lacking promoter), or V3 (derived from V1 with modified primer binding sites). All dark grey bars are at the limit of detection, where a pseudocount of 1 was used for calculating detection limits.

The initial transposon donor (V1) carried a selectable *acrVIA1* cassette flanked by a ΦKZ consensus early promoter^8,26^ and a terminator, to drive early *acrVIA1* expression (Table S1). For selection, we co-expressed *Listeria seeligeri* Cas13a (LseCas13a)^19,27^ with two crRNAs: one targeting *orf120* (major capsid) and one targeting *orf55* (a nvRNAP subunit). Each crRNA alone strongly restricted phage infection but was individually escapable (Figure 1C). We therefore designed a single repeat–spacer–repeat–spacer array expressing both crRNAs, which efficiently blocked infections by previously isolated escaper phages (Figure S1A). No escaper mutants were detected (limit of detection (LOD) <2.5 x 10^-10^ pfu/mL), so this dual crRNA array was used for all subsequent selections (Figure 1C, Figure S1B).

To generate a transposed phage population, we infected a log-phase culture expressing Mariner–Nlp1 with ΦKZ for 16 h and then selected output phages on the dual-crRNA Cas13a strain using full-plate infections. Single plaques were isolated, and *acrVIA1* was detected by PCR (Figure S1C). Plaque formation required both the transposase and the transposon donor (Figure 1D–E, Figure S1D). Whole-genome sequencing of four independently isolated phages confirmed bona fide transposon insertions in *orf303*, *orf208*, *orf241*, and *orf252*. In each case, two inverted repeats flanked an insertion at a TA dinucleotide in the ΦKZ genome (schematized in Figure S1C).

Transposition using the V1 donor occurred at a rate of ∼1 in 3 × 10^7^ phages, demonstrating that transposition is inefficient but feasible in a large population (Figure 1E). No plaques were detected when the donor plasmid was present without transposase (LOD: 1.3 x 10^-10^ pfu/mL (Figure 1E)). A second donor (V2) lacking the early promoter yielded slightly lower efficiency (1 in 6.7 × 10^7^ phages; Figure 1E), consistent with *acrVIA1* expression relying on endogenous promoters. To facilitate compatibility with bacterial Tn libraries, we engineered a third donor (V3, derived from V1) that altered the transposon inverted repeat junctions to introduce orthogonal primer binding sites. Transposition efficiency with V3 (1 in 7 × 10^7^ phages) was similar to V1 and V2 and remained strictly dependent on the Nlp1–Mariner fusion (LOD: 3.9 x 10^-11^ pfu/mL). Transposition was detected across a range of input phage concentrations and was most efficient at a relatively low input (∼2 × 10^4^ pfu/mL; Figure S1E). We therefore used the V3 donor to generate a pool of mutants for insertion sequencing.

### A ΦKZ phage TnSeq library with genome-wide coverage

To build independent pools of transposon mutants, we performed ten biological replicate transposition infections (Figure S2). Mutants from each reaction were enriched on the Cas13a counter-selection host, and fractions from all ten pools were combined to generate a single mutant library. We isolated DNA, prepared sequencing libraries, and amplified transposon–genome junctions for Illumina sequencing (Methods). We obtained ∼3.4 million high-quality reads, 90% of which contained transposon sequence. Of these, 55.7% mapped to the ΦKZ genome and the remainder to the V3 donor plasmid. Across the ΦKZ genome, 6,667 unique insertion sites were identified (Figure 2A), corresponding to a mean of one insertion every 42 bp over the 280,334 bp genome.

**Figure 2.**
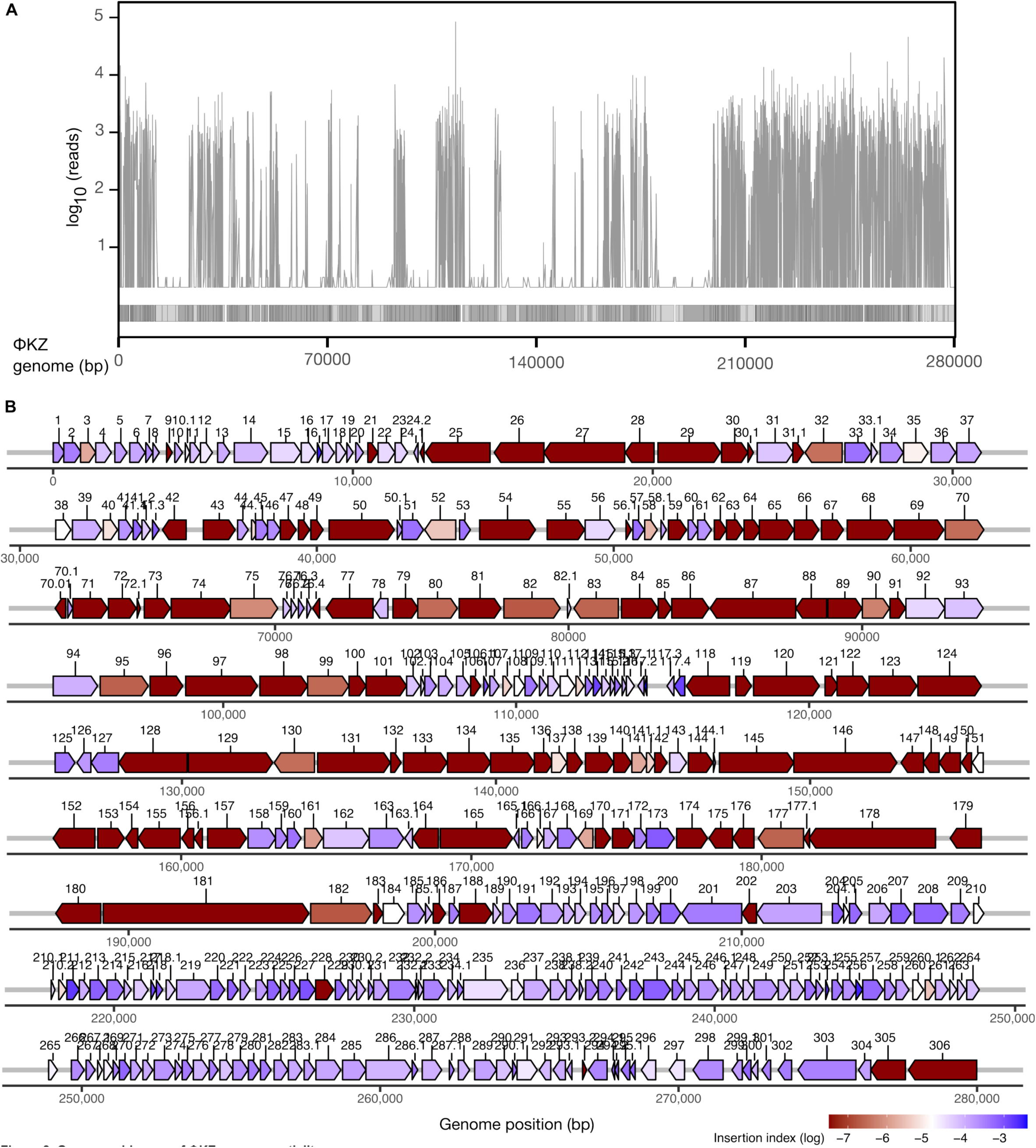
Genome-wide map of ΦKZ gene essentiality. (A) Genome-wide distribution of transposon (Tn) insertion reads across the ΦKZ genome. All forward reads are shown without filtering, with a pseudocount added to sites with single reads for log-scale visualization. The ΦKZ genome map is displayed below, with genes in the forward direction in dark gray and reverse in light gray. (B) Schematic summary of gene essentiality across the ΦKZ genome, as determined by insertion sequencing (TnSeq) analysis.

To define confident gene insertions, we first examined genes expected to be essential. Many insertion sites yielded low sequencing depth, suggesting that insertion occurred but rendered phages unfit. We therefore required >4 reads per insertion for downstream analysis. In addition, insertions within the first or last 10% of a gene were excluded to avoid misclassifying overlapping genes or insertions that might not fully disrupt gene function. Using these criteria, ∼30% of ΦKZ genes lacked disruptive insertions, whereas ∼70% tolerated insertions (Figure S3).

### ΦKZ phage TnSeq identifies essential genes

As an initial accuracy check, we focused on large genes, which should, by chance, accumulate many insertions unless they are essential. Tailed phage genomes often encode large essential structural and polymerase proteins. In our ΦKZ TnSeq dataset, all 16 genes >2,154 bp and 30 of 34 genes >1,560 bp lacked disruptive insertions (Figure 2B), strongly indicating essentiality given the genome-wide insertion frequency (one unique insertion every 42 bp). Below, we highlight examples of essential genes in tail, capsid, and conserved non-virion modules with all features reported in Figure 3A

**Figure 3.**
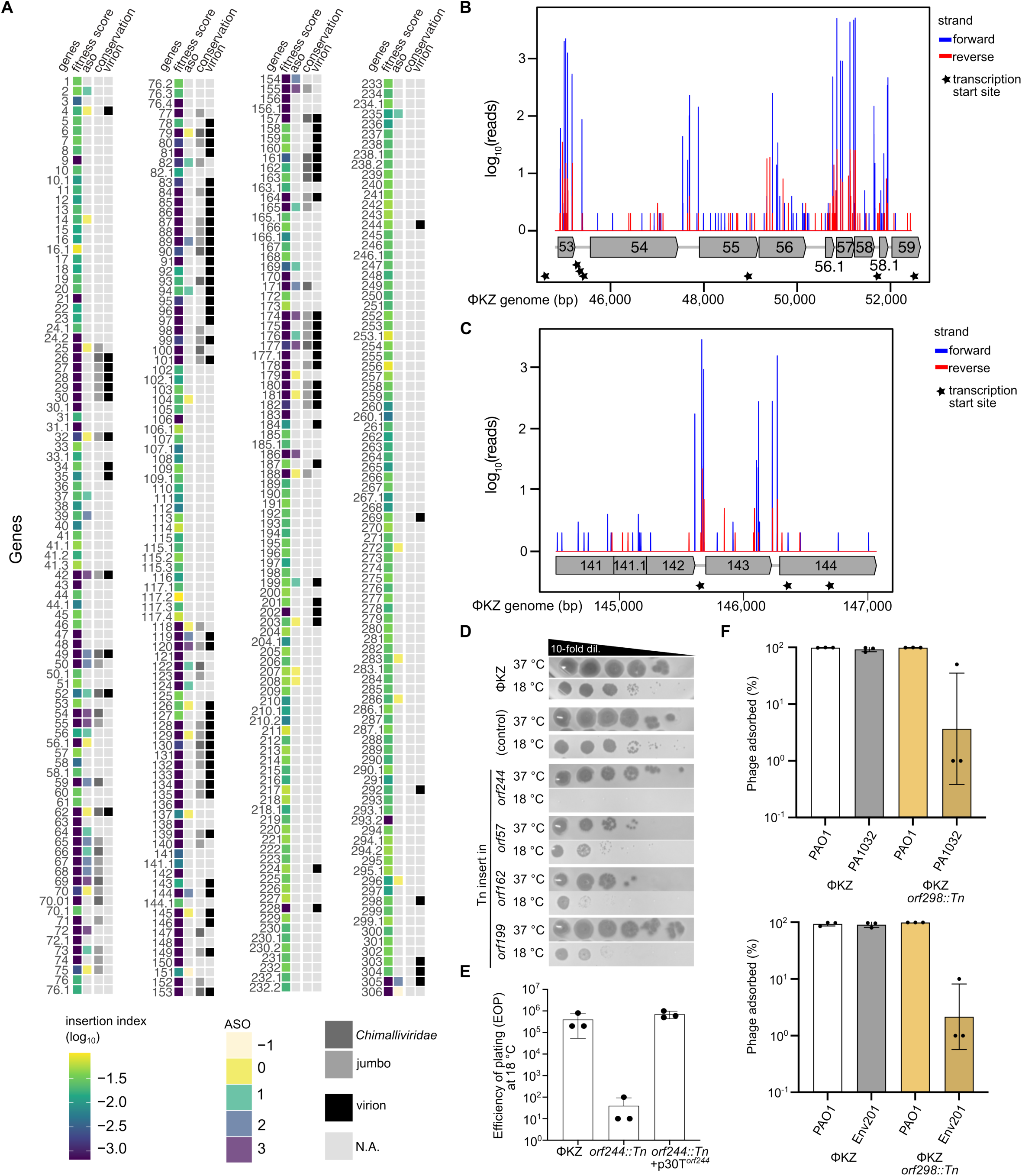
Discovery of novel ΦKZ gene functions using Tn mutagenesis. (A) Heat map showing the insertion index derived from phage TnSeq. Yellow indicates many tolerated insertions, while darker shades indicate fewer insertions. ASO: Fitness scores after ASO knockdown^20^, with yellow (scores −1, 0) indicating no phage fitness cost detected after knockdown while darker shades (1, 2, 3) indicate increasing fitness costs upon gene knockdown. Conservation: Genes conserved across related jumbo phages (jumbo) are indicated as well as the subset of conserved genes that are uniquely found amongst *Chimalliviridae*. Virion: the protein was detected in the virion. (B–C) Examples of Tn insertion read density across representative genomic regions (*orf 53–59* and *141–144*). (D) Plaque assays showing environmental temperature-sensitive mutants at 37 °C and 18 °C. Wild-type ΦKZ and a transposed control phage are included. (E) Adsorption efficiency of wild-type and *orf298::Tn* ΦKZ phages in *P. aeruginosa* PAO1 and clinical isolates.

#### Tail genes

The largest gene in ΦKZ, *orf181*, encodes a lysozyme-containing tail protein likely serving as the tape measure^28^. *orf181* and its neighbor *orf182* (encoding a baseplate protein) both lack transposon insertions (Figure 2B). Given the 6,714 bp size of *orf181*, its complete lack of insertions makes it a high-confidence essential gene. Other tail-associated genes lacking Tn insertions include putative tail fibers (gp145, gp146), a putative baseplate wedge protein (gp87), a paralogous baseplate protein family (gp131–135)^28^, the tail sheath^29,30^ (gp29), and the tail tube (gp30) (Figure 2B). Virion-associated proteins of unknown function such as gp27 (DUF7193) and a low-copy virion protein gp128 also lack insertions. Virion localization and functional annotations were supported by previously published mass spectrometry datasets^31–33^ and HHpred/PSI-BLAST analyses (Figure 3A).

#### Head genes

Several large genes that appear essential by phage TnSeq encode for capsid assembly, DNA packaging, or injected proteins. These include the major capsid protein (gp120)^32,34^, the portal (gp129)^32,35^, and the large terminase (gp25)^32^ (Figure 2B). Mass spectrometry of a tail-less ΦKZ mutant revealed >50 distinct polypeptides incorporated into the head, many of which are proteolytically processed by an essential (per phage TnSeq) head protease, gp175^32,36^. Many head proteins (gp92, 93, 94, 95, 96, 97, 153, 162, 163; PF12699.12 (C-terminus)^10,32^) are members of a paralogous family^10,33^ and some of them are injected during infection^10^. Interestingly, several paralogous genes (e.g., *orf92, 93, 96, 162, 163*) readily tolerate insertions, whereas genes encoding other high-copy inner body proteins^32^ such as *orf89* and *orf90*, along with genes encoding the vRNAP (*orf178, 149, 180, 80*)^8,37^ appear essential by phage TnSeq.

To validate a subset of phage TnSeq-based essentiality calls, we used Cas13a to target transcripts encoding head proteins and isolated escape mutants. Strong targeting yielded phages with large deletions in non-essential genes or point mutations in essential genes. Deletions were obtained in several genes predicted to be non-essential by Phage TnSeq (*orf93, 94, 162*, *163, 203, 303*), while targeting of genes predicted to be essential (*orf89*, *90*, *99*, *119*, *177*) did not result in deletions (Figure S4A-C), corroborating their essentiality. Notably, *orf97*, *orf157*, and *orf85*–*86* tolerated deletions despite lacking transposon insertions, likely due to polar effects resulting from insertions (described below). *orf95* is dispensable by Cas13a targeting but incurs a fitness cost in competition (Figure S5).

#### Conserved *Chimalliviridae* Proteins

In addition to gene size, evolutionary conservation is expected to be correlated strongly with essentiality. A previous study defined 20 ΦKZ genes that are conserved across Chimalliviridae yet unique to this phage family^38^. Of the “Prichard 20”, 19 appear essential or strongly fitness conferring by phage TnSeq (15 with zero insertions; 4 with only 1–3 insertions; Figure 3A, Supplementary Data 1). The lone exception is the paralogous protein family encoded in multiple copies (*orf93/162/163*). As an example of an essential conserved gene, Chimallin A (*orf54*), the phage nucleus shell protein^39,40^, was previously shown to be refractory to knockout in ΦKZ^19^ and essential upon translational repression in the related *E. coli* phage Goslar^9^. No Tn insertions were identified in this gene. Other conserved genes encoding protein importers (*imp1*/*orf69*; *imp2*/*orf47*; *imp3*/*orf59*; *imp4*/*orf48*), non-virion RNA polymerase (nvRNAP) subunits (*orf123, orf74, orf68, orf71–73, orf55–56.1*), and the nucleus-associated *imp6/chmC/orf67*^25,41^ also lack disruptive insertions (Figures 2B, 3A). Additional conserved non-virion genes, such as *orf165* (a putative SbcCD-like ABC ATPase with SMC homology), also lacked insertions. A subset of these genes were also queried for essentiality previously via ASO-based translational repression^20^, with generally consistent results and some exceptions. Figure 3A summarizes the concordance between phage TnSeq fitness scores and ASO knockdown.

### Polar Effects and Transcriptional Architecture

Insertional approaches in bacteria and phages are susceptible to polar effects, where an insertion disrupts expression of downstream genes in the same transcriptional unit. Polar effects are more severe when a small number of operons are driven by few transcription start sites (TSSs), and less problematic when multiple TSSs distribute transcription across densely packed loci. Although phages are sometimes depicted as having few promoters, recent short-read RNA sequencing identified 149 TSSs across the ΦKZ genome^26^, in line with earlier predictions of 134 operons^8^.

To ask whether abundant TSSs mitigate polar effects in ΦKZ, we again examined the *chmA* (*orf54*) gene. *orf53* and *chmA* appear in a putative operon with a TSS driving *orf53-chmA*, yet multiple transposon insertions are present in *orf53* while none occur in *chmA* (Figure 3B). This indicates that Tn insertion in *orf53* does not abrogate *chmA* transcription. An independent TSS is found directly upstream of *chmA,* likely sustaining *chmA* transcription even when *orf53* is disrupted. *orf55* (nvRNAP subunit) also appears essential, while *orf56–57* are non-essential and *orf58–59* appear essential (Figure 3B). A TSS upstream of *orf59* (*imp3*) supports the interpretation that the absence of insertions in *orf58* reflects essentiality of gp58, rather than a polar effect on *imp3*. This pattern in a tightly packed locus underscores how multiple TSSs can insulate neighboring essential genes from polar effects.

Another example of polar effects being mitigated arises in the lysis cassette. Biochemical studies show that gp144 is a highly active peptidoglycan hydrolase and the likely endolysin^42,43^. A recent preprint further identified gp142 as the holin and gp143 as a lysis regulator^44^. By phage TnSeq, *orf142* and *orf144* lack insertions and appear essential, whereas *orf143* is disrupted by five insertions in its 534 bp coding sequence. Multiple TSSs in this region are again identified, upstream of *orf139*, *orf143*, and *orf144*^26^ (Figure 3C). Thus, many potential polar effects are mitigated by the underlying transcriptional architecture. Importantly, abundant TSSs are not unique to ΦKZ: several smaller *Pseudomonas* phages profiled by the same method also exhibit numerous TSSs (e.g., 14–1, 50 TSSs; LUZ24, 16; LUZ19, 9; PAK_P3, 75; YuA, 21)^26^.

By contrast, polar effects are expected when genes are expressed from a single polycistronic transcript without internal TSSs. Two clear examples are the ΦKZ intron-encoded homing endonucleases gp72 and gp179, both lying within RNA polymerase operons. ASO knockdown data suggested that gp72 is essential while gp179 is not^20^, while both genes appear essential by phage TnSeq (no insertions). Moreover, a gp179 homolog in ΦPA3 is non-essential but implicated in phage–phage competition^45^. To resolve this discrepancy, we used Cas13a to target each gene and readily isolated deletion mutants in *orf72* and *orf179*, demonstrating that neither gene is strictly essential for ΦKZ replication (Figure S4C). These loci thus exemplify polar effects: transposon insertion in *orf72* or *orf179* likely terminates transcriptional readthrough, blocking expression of the downstream essential genes (*orf73* or *orf180*). Similar cases likely exist elsewhere in the ΦKZ genome. Cas13-based targeting of individual genes coupled to sequencing of escape mutants provides a straightforward way to discriminate true essential genes from polar effect artifacts uncovered by phage TnSeq.

### Non-essential genes revealed by dense insertion coverage

Genes with abundant, evenly distributed insertions can be confidently classified as non-essential. *orf39* encodes the PhuZ tubulin required for nucleus positioning^12,13,46,47^ contains many insertions across its length and has been previously deleted^19^. ChmB (gp2), a protein proposed to cooperate with PhuZ to facilitate capsid treadmilling and docking at the phage nucleus^48^, likewise appears non-essential, suggesting that core aspects of DNA packaging remain to be fully explained.

Other genes with many Tn insertions and known non-essential functions encode: a ribosome binding protein (gp14)^49^, an injected protein^19^ (gp93), Thoeris anti-defense protein 1^50^ (Tad1, gp184), a cluster of uncharacterized non-essential proteins gp206-216^19^, the activator of Juk defense (gp241)^51^, and proteins encoded nearby (gp237-242)^51^. Dip (gp37), a previously studied RNA degradosome inhibitor^52^, also appears dispensable. Interestingly, most virion proteins in ΦKZ appear essential, but notable exceptions include a helicase gp203 and gp303 (unknown function), both of which are proteolytically processed and packaged into the head^32^. gp203 and gp93 co-pellet with phage genomic DNA after capsid disruption^32^, indicating tight association with packaged DNA despite being non-essential under the conditions tested Small genes were particularly likely to tolerate insertions. Of 210 genes shorter than 500 bp (arbitrarily defined as “small”), 185 contain at least one insertion, while just 25 lack insertions. Because small genes are more likely to be misclassified as essential by chance, this likely underestimates the true number of non-essential small genes. Even so, our insertion data strongly indicate that approximately half of ΦKZ genes are both small and non-essential under the laboratory conditions used here (37 °C, LB, aeration). Across the full genome, roughly 70% of genes appear non-essential under these conditions. Many of these non-essential genes may be conditionally essential in specific hosts or environments (examples are presented in the next section)—phenotypes that phage Tn mutagenesis is uniquely poised to uncover.

### Minor capsid proteins are required at environmental temperature

*P. aeruginosa* is frequently isolated from soil, freshwater, and human-associated environments^53,54^. We therefore queried whether any phage genes are dispensable at 37 °C but required at a lower environmental temperature (18 °C). We subjected a randomly chosen set of 90 transposed mutants to infection at 18 °C. Four mutants that grew poorly at 18 °C were identified and sequenced alongside a control mutant that grew normally (Figure 3D). Insertions mapped to *orf162*, *orf199* (two redundant insertions), and *orf244*. All three genes encode proteins packaged in the capsid^32^. gp244 is notable as it appears to decorate the outside of the phage capsid^34^. The *orf244::Tn* mutant completely failed to replicate at 18 °C, a defect rescued by expressing *orf244*^33^ in trans (Figure 3E).

To understand the role of minor capsid proteins in this phage, we next checked what phage TnSeq and a recent cryo-EM structure of the capsid reveal^34^. The cryo-EM structure revealed ∼10 “minor capsid” proteins whose N-termini weave into major capsid hexamers^34^. Most of the genes encoding these minor capsid proteins, including *orf35*, *orf93*, *orf162*, *orf184* (*tad1*), and *orf244*, appear non-essential by phage TnSeq, at 37 °C. Interestingly, only three minor capsid genes appear essential at 37 °C: *orf28*, *orf91*, and *orf119*. The structural data provide an explanation for this, revealing a junction between vertex-binding complexes in which gp28 from one complex contacts gp91 and gp119 from the adjacent complex^34^. Cas13a targeting of these genes generated escapers lacking deletions, confirming essentiality and supporting a model where they form an interlocking complex critical for capsid assembly (Figure S4B). These findings therefore suggest multiple roles for minor capsid proteins: i) three minor capsid proteins that form an essential (at 37 °C) complex in the capsid, ii) additional minor capsid proteins, including a capsid decorating protein gp244, that are dispensable at 37 °C, but essential at 18 °C, and iii) non-essential proteins whose N-termini are wedged into the capsid and the C-termini that are liberated by a head protease^32^ are injected during the next round of infection (i.e. gp93, gp184/Tad1)^33,50^. The transposed mutant library and pooled TnSeq can thus be used to rapidly identify genes whose contributions emerge only in ecologically relevant conditions—a likely major evolutionary pressure for *P. aeruginosa* phages^55^.

### gp298 is a conditionally essential tail protein for adsorption to wild isolates

Phage Tn mutagenesis provides a straightforward route to isolate knockouts for every non-essential gene, enabling systematic screens for strain- or condition-specific phenotypes. To explore host-range determinants, we arrayed ∼1,500 transposed ΦKZ mutants in 96-well format and plated the collection on 18 wild *P. aeruginosa* strains. Host-specific phage “dropouts” were readily detected (Figure S6).

One striking example is *orf298*, which has 28 insertions across its 1,008 bp coding region after propagation in PAO1, yet an *orf298::Tn* mutant failed to plaque on clinical isolate PA1032 (from a human acute infection, San Francisco) and on Env201 (a soil isolate from Toronto). gp298 contains a Kelch domain and a predicted N-acetylneuraminate epimerase domain and is present at low abundance in the virion^33^. In infection time courses, wild-type ΦKZ efficiently adsorbed to all three hosts (as indicated by decreased free phage titers within 15 min), whereas the *orf298::Tn* mutant specifically failed to adsorb to PA1032 and Env201 but adsorbed normally to PAO1 (Figure 3F). These data support gp298 as a previously unrecognized tail-associated factor required for adsorption and/or receptor access on a subset of wild *P. aeruginosa* strains. Notably, *orf298* is not located near canonical tail genes; instead, it clusters with other non-essential genes of unknown function.

### Nlp1 is a fitness-conferring recombinase required for NucC resistance

To identify phage genes that are conditionally essential for DNA replication or genome stability, we screened the 1,500 arrayed ΦKZ mutants against two nucleases: EcoRI and the Type III-C CBASS system with a NucC effector. Wild-type ΦKZ naturally resists both^14,25^. We hypothesized that mutants with increased nuclear permeability might become sensitive to EcoRI, a ∼30 kDa protein that cannot normally access the phage genome but cleaves ΦKZ DNA if granted access to it *in vivo* and *in vitro*^14^. However, none of the mutants were restricted by EcoRI, indicating that genomic segregation from the cytoplasm—and thus resistance to both EcoRI and the endogenous PAO1 Type I RM system^14,56^—is robustly maintained across all tested mutants.

By contrast, one ΦKZ mutant completely lost replication on strains expressing a Type III-C CBASS system (Figure 4A). This system senses unknown phage factors, produces cAAA signals, and activates NucC, a nuclease effector previously implicated in blocking a ΦKZ-like phage in *Serratia*^57^. In that context, degradation of the host genome outside the phage nucleus was proposed to halt phage progression. We were therefore particularly interested in mutants that become sensitive to NucC, because ΦKZ-like phages degrade the host genome^46,47^.

**Figure 4.**
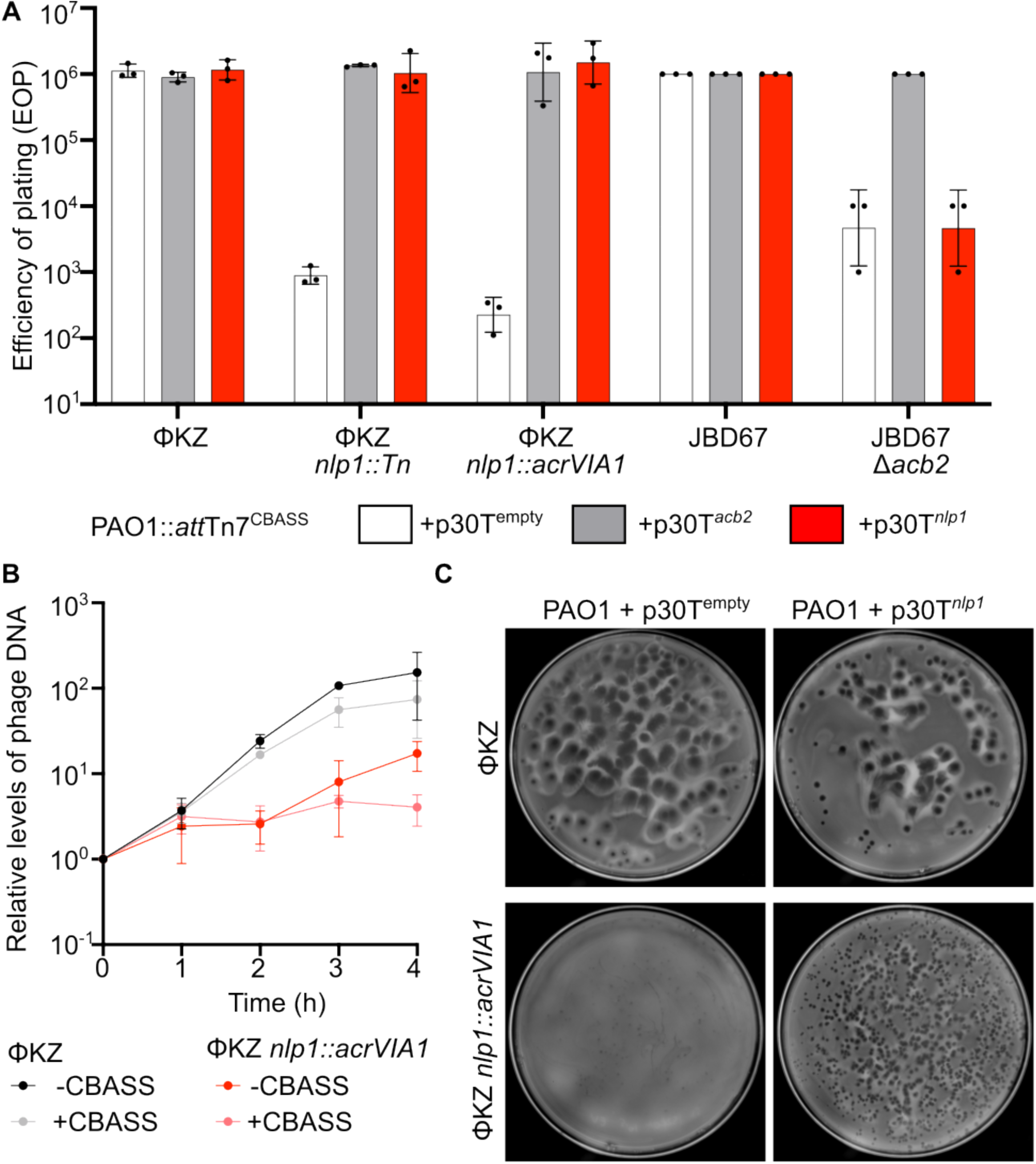
Nlp1, a ΦKZ-specific UvsX homolog, protects against CBASS defense. (A) Efficiency of plating (EOP) of wild-type and Δ*nlp1* ΦKZ in the presence of CBASS. Control phages JBD67 and JBD67Δ*acb2* confirm CBASS targeting specificity, while Acb2 expression neutralizes CBASS activity. (B) Quantitative PCR analysis of ΦKZ wild-type and ΦKZ *nlp1::acr* ΦKZ and replication in the presence and absence of CBASS. (C) Full-plate infections comparing ΦKZ and ΦKZ *nlp1::acrVIA1* with or without *nlp1* complementation.

The NucC-sensitive mutant carried a Tn insertion in *nlp1* (nucleus-localized protein 1), encoding a protein homologous to phage T4 UvsX^58^. UvsX is a recombinase that is not strictly essential in T4, but its loss confers sensitivity to DNA damage^58^. To confirm this result, we deleted *nlp1* by replacing it with *acrVIA1* (ΦKZ *nlp1::acrVIA1*, hereafter referred to as Δ*nlp1)* under complementation with wild-type *nlp1 in trans*. Like the Tn mutant, the Δ*nlp1* phage was sensitive to Type III-C CBASS NucC, and this sensitivity was rescued by Nlp1 expression in trans (Figure 4A). Sensitivity was dependent on canonical CBASS activity because Acb2, a cyclic-nucleotide sponge that sequesters cAAA^59^, rescued the Δ*nlp1* phage. In contrast, Nlp1 expression did not rescue CBASS NucC sensitivity of an unrelated phage (JBD67Δ*acb2*), indicating that Nlp1 is not a general CBASS inhibitor (Figure 4A).

During infection, NucC activity modestly reduced WT ΦKZ DNA levels yet completely abolished DNA replication in the Δ*nlp1* mutant (Figure 4B). Attempts to visualize NucC localization were unsuccessful because fluorescent tagging rendered NucC inactive. Nonetheless, the genetic data support a model in which Nlp1’s recombinase activity repairs NucC-induced phage DNA damage, analogous to UvsX-mediated recombination of fragmented T4 genomes after UV or Cas9/Cas12 cleavage^60^. While NucC is often viewed as anti-phage through host genome destruction, our data suggest that NucC can directly damage phage DNA and that ΦKZ relies on Nlp1 to tolerate this assault. Consistent with a DNA repair defect specific to NucC, Δ*nlp1* phage remains fully resistant to EcoRI (Figure S7).

The absence of Tn insertions in *nlp1* in the pooled phage TnSeq library suggested that it is a strongly fitness-conferring gene. Δ*nlp1* phages replicated slower (Figure 4B), formed smaller plaques than wild-type ΦKZ, and plaque size was restored by Nlp1 complementation (Figure 4C), a phenotype reminiscent of T4 *uvsX* mutants^58^. Tn insertions in *nlp1* were likely present at low frequency in the initial transposed population but were outcompeted during propagation. In a direct competition assay over 4 h in liquid culture, control acr insertions in *orf2/chmB*, *orf184/tad1*, *orf39/phuZ*, or *orf241* exhibited competitive indices of ∼0.35–2.0 (Figure S5), while Δ*nlp1* showed a competitive index of 0.05, confirming a strong fitness defect. *nlp1* thus exemplifies a gene for which full deletion is possible, but the mutant drops out during phage competition or in the presence of CBASS NucC.

Together, these findings highlight the value of pairing pooled phage TnSeq to map essential/fitness genes with an arrayed mutant library to interrogate conditional phenotypes and recover mutants that are underrepresented in pooled selections.

### Non-essential gene clusters and defense-related functions

Many non-essential ΦKZ genes cluster within a broad region of largely contiguous non-essential genes spanning from ∼*orf184* through *orf304* and then *orf1*–*orf20* on the left arm (Figure 2B). ΦKZ is circularly permuted and its genome ends are arbitrary. This region roughly corresponds to the portion absent from a reduced size *P. aeruginosa* jumbo phage, ΦEL^35,61^. This observation suggests that this region encodes host-interacting or defense-related adaptations rather than core replication modules.

It is tempting to speculate that these non-essential regions constitute “anti-defense loci”, as described for smaller phages^62^, although this remains to be systematically tested. Our arrayed mutant screens offer initial support. When we plated the 1,500 mutants on *P. aeruginosa* strain Env201, we identified a mutant with an insertion in *orf302* that failed to form plaques. This phage adsorbed normally but failed to produce progeny, consistent with an Env201-specific intracellular defense that is absent in PAO1 (Figure S8). In a separate screen against the PfJuk^51^ (jumbo phage killer) defense system, which restricts ΦKZ, we observed robust restriction of most mutants but identified one that grew well. Sequencing revealed a Tn insertion in *orf241*, encoding the previously described Juk activator^51^. Thus, the transposed mutant collection can rapidly uncover both inhibitors and activators of diverse defense systems across wild *P. aeruginosa* isolates.

## Discussion

Phage functional genomics is beginning to match the scale and sophistication of bacterial genetics, but tools remain limited, particularly for large and complex phage genomes. Here, we establish a generalizable phage transposon mutagenesis and phage TnSeq platform that (i) maps essential and fitness-conferring genes genome-wide, (ii) generates a library of knockouts in nearly all non-essential genes, and (iii) identifies safe harbor loci suitable for insertion and expression of transgenes.

By directing *mariner* transposition into the ΦKZ nucleus and selecting for insertions using an anti-CRISPR and CRISPR-Cas13a, we demonstrate a rapid and scalable workflow for phage genome mutagenesis. Because *mariner* is widely used in bacteria, requires no host-specific factors, and inserts semi-randomly, we anticipate that this approach will be broadly adaptable across bacteria–phage pairs. The transposon donor can be easily reconfigured—for example, by removing or modifying promoters—to profile therapeutic phages and identify optimal genomic contexts for stable expression of payload genes such as anti-defense factors or therapeutic cargo.

Phage TnSeq is highly complementary to recently developed ASO and CRISPRi-ART platforms^20,21^, which are particularly suited to titrating expression of essential genes. In contrast, TnSeq requires no guide RNA or oligo design, can be deployed at genome scale in a single experiment, and yields hundreds to thousands of individual insertion mutants that can be archived and screened under diverse conditions. Tn insertions, however, are not ideal for fine-tuned modulation of essential gene expression and are inherently limited by issues such as polar effects. Moving forward, we anticipate that the recent explosion in defense and anti-defense system discovery means that one can have the ability to transpose, and select for transposed plaque formation, with any phage of interest.

One of the most striking outcomes of this work is the revelation that a majority of ΦKZ genes are dispensable under standard laboratory conditions. With >200 dispensable genes, it is surprising that gp184/Tad1 was the only previously known anti-defense gene in ΦKZ^50^. Some of these non-essential genes encode tail (e.g. gp298) or capsid-associated proteins (e.g. gp244) with host- or temperature-specific roles, defense inhibitors (e.g. gp302) or activators (e.g. gp241), and a recombinase (Nlp1/gp152) that becomes essential in the presence of a specific nuclease. The ease with which transposon mutagenesis can interrogate this abundant pool of non-essential genes sets the stage for systematic dissection of phage accessory and anti-defense functions.

Smaller phages likely encode a lower fraction of non-essential genes than ΦKZ, yet the total number of non-essential genes per phage may still be modest enough that a complete collection of knockout mutants could be arrayed in a single or a few 96-well plates. This makes it feasible to screen for defense inhibitors, host-range factors, and conditionally essential genes across multiple hosts, temperatures, and selective pressures. As interest grows in phage diversity, basic biology, genome engineering, and therapeutic applications, we anticipate that phage Tn mutagenesis will provide a broadly useful resource and methodology. The essentiality maps, mutant libraries, and design principles described here should be transferable to other phage systems, enabling the community to move from sequence catalogs to functional insight at scale.

### Limitations

Phage TnSeq, as implemented here, requires a robust defense system that is difficult for phages to escape and a cognate anti-defense factor that can be encoded on the transposon. These requirements may constrain applicability for some phage–host systems. Nevertheless, the programmability of CRISPR-Cas systems and the growing catalog of anti-CRISPR proteins^63^ provide a flexible toolbox for adapting selection schemes.

A second, inherent limitation of insertional mutagenesis in prokaryotes and phages is polar effects: insertions can appear to reveal essential genes when they instead disrupt expression of neighboring essential genes. We show that the high density of TSSs in ΦKZ alleviates many potential polar effects, but we also identify clear cases where phage TnSeq misclassifies non-essential homing endonucleases as essential due to operon structure. In some regions, we identified insertion direction biases suggesting that perhaps read through transcription can alleviate or induce confounders. Using Tn donors that lack transcriptional terminators or that incorporate unidirectional terminators could reduce some polar effects. Consequently, putative essential genes of interest should be addressed by complementary approaches such as targeted knockdowns or targeted deletions.

## Acknowledgements

J.B.-D. is supported by the National Institutes of Health (nos. R01 AI171041 and R01 AI167412). C.K. received support from the UCSF Discovery Fellowship and the NIH award 2T32AI060537-21. D.M. received support from the NIH Ruth L. Kirschstein National Research Service (NRSA) award 1F32GM149125-01/-02. We thank Bondy-Denomy laboratory members for input into this work.

## Author contributions

A.W.C. - Conceptualization, Formal analysis, Investigation, Methodology, Validation, Visualization, Software, Writing - original draft,

W.X.Y. - Investigation, Methodology, Validation, Visualization, Writing - original draft, Writing review and editing, Writing - original draft, Writing - review and editing

D.M. - Investigation, Visualization, Writing - original draft, Writing - review and editing

C.K. - Investigation, Visualization, Writing - original draft, Writing - review and editing

M.R.M. - Investigation

Y.L. - Conceptualization, Resources

Y.Y. - Resources

J.B.D. - Conceptualization, Funding acquisition, Investigation, Writing - original draft, Writing review and editing

## Supplemental Table Legends

**Table S1. List of oligonucleotides used in this paper**

**Table S2. Raw TraDIS output for gene essentiality after trimming**

**Table S3. Raw TraDIS output for all insertions across the ΦKZ genome**

## RESOURCE AVAILABILITY

### Lead contact

Further information and requests for resources and reagents should be directed to and will be fulfilled by the lead contact, **Joseph Bondy-Denomy**.

### Materials availability

Plasmids and strains generated in this study are available from the lead contact upon reasonable request and, where possible, will be deposited with a public repository (e.g., Addgene).

### Data and code availability

- Insertion sequencing data and ΦKZ whole-genome sequencing data will be deposited in a public repository and accession numbers will be provided upon publication.
- Custom analysis scripts are available from the lead contact upon request.

## EXPERIMENTAL MODEL AND SUBJECT DETAILS

### Bacterial strains

Unless otherwise indicated, experiments were performed in *Pseudomonas aeruginosa* PAO1 and PAO1 derivatives. Bacteria were grown in LB broth at 37 °C with aeration. Where appropriate, media were supplemented with gentamicin (50 μg/mL) or carbenicillin (250 μg/mL).

### Phages

The jumbo phage ΦKZ and derivative mutants generated in this study were propagated on *P. aeruginosa* PAO1 or specified host strains. For host-range and conditional screens, a panel of wild *P. aeruginosa* isolates, including clinical isolate PA1032 and environmental isolate Env201, were used as indicated.

## METHOD DETAILS

### Construction of mariner transposase expression strains

The mariner transposase gene was amplified from pBTK30 (kindly provided by the Lory lab) and assembled into a Tn7 plasmid carrying sfCherry–gp152. For construction of FLAG-tagged transposase, plasmid p30T expressing sfCherry-Mariner–Nlp1 was linearized with primers oAWC088/oAWC021, and the resulting backbone was ligated with gBlock oAWC093 using NEBuilder HiFi DNA Assembly (New England Biolabs).

All plasmids were sequence-verified (Quintara Biosciences) and introduced into *P. aeruginosa* PAO1 by electroporation together with pTNS3 for integration at the Tn7 site^1^. Transformants were selected on LB agar containing 50 μg/mL gentamicin. The Flp recombinase plasmid pFLP was then introduced into these strains to excise the gentamicin resistance cassette. Colonies were counter-selected on sucrose, and loss of gentamicin resistance was confirmed by replica plating.

### Construction of *acrVIA1* transposon donors (V1–V3)

For construction of *acrVIA1* transposon donors V1 and V2, p30HERDT (abbreviated p30T) was linearized with primers oAWC047/oAWC048. gBlocks encoding distinct constructs (oAWC122 and oAWC123 for donors V1 and V2, respectively) were ligated into the linearized backbone using HiFi Assembly and transformed into *E. coli* XL1-Blue.

For the V3 donor used in pooled TnSeq experiments, the p20T plasmid was linearized instead of p30T and ligated with gBlocks oAWC146 and oAWC147 using HiFi Assembly. All constructs were confirmed by Sanger sequencing and subsequently transformed into PAO1 strains carrying the mariner transposase, as described above.

### Construction of pHERD30T expression plasmids

To generate gene inserts in p30T, the plasmid backbone was first amplified with primers WX_016 and WX_017. Genes of interest were amplified with the appropriate primer pairs and assembled into the backbone using HiFi Assembly. Constructs were cloned into XL1-Blue, sequence-verified, and introduced into PAO1 by electroporation.

### Construction of CRISPR-Cas13 guide plasmids

For cloning of Cas13 guide arrays for single or dual targeting, the plasmid backbone was amplified with primers oAWC004/oAWC024. Complementary oligonucleotides (Table S1) encoding the desired crRNAs (100 μM each) were annealed in Duplex Buffer A and ligated into the backbone using HiFi Assembly.

For Cas13 guide plasmids used to target candidate essential or non-essential genes in ΦKZ (to isolate escaper mutants), the Cas13 backbone plasmid was amplified with primers that incorporated the crRNA sequence directly. Products were assembled with HiFi and cloned into XL1-Blue. All guide plasmids were confirmed by sequencing and transformed into PAO1 or PAO1 *attTn7::cas13aLse* as appropriate.

### Generation of transposed ΦKZ arrays

Transposed ΦKZ phages were selected on a Cas13 strain expressing dual crRNAs targeting *orf120* and *orf55*. Individual plaques were picked into SM buffer. In total, 1,800 plaques were collected from 20 plates in a 96-well format, with plate corners reserved for wild-type ΦKZ controls and two central wells containing no phage. The phage array was replicated onto lawns of different *P. aeruginosa* isolates and incubated overnight at 37 °C to assess host-range and conditional phenotypes.

### Cas13a-mediated isolation of ΦKZ deletion mutants

High-titer ΦKZ stocks were plated on induced PAO1 *attTn7::cas13aLse* strains carrying targeting guides as described above. Escaper plaques were picked and propagated on PAO1 *attTn7::cas13aLse* expressing the same guides. Regions spanning the putative deletion were amplified using primers flanking the targeted locus and sequenced to confirm the nature of the deletions or point mutations.

### Deletion of individual ΦKZ genes by *acrVIA1* replacement

Homology-directed constructs for gene replacement with *acrVIA1* were synthesized (Twist Bioscience) with compatible overhangs and cloned into NheI-digested p30T using NEB HiFi Gibson assembly. Constructs were selected with gentamicin and confirmed by sequencing.

Phage recombinants were generated and selected similarly to a previous report^2^. PAO1 strains harboring these plasmids were grown overnight, subcultured 1:100 into fresh LB, and grown at 37 °C to an OD₆₀₀ of 0.6. Cultures were diluted 1:100, and ∼10³ PFU of ΦKZ were added for overnight infection. The following day, cultures were centrifuged at 10,000 × g for 2 min, and chloroform was added to the supernatant at a 1:8 (v/v) ratio. After ≥15 min at room temperature, lysates were centrifuged again to clarify.

For selection of recombinants, 50 μL of lysate were mixed with 300 μL of PAO1 *attTn7::cas13aLse* + p30T crRNA^55/120^ in the presence of 0.3% arabinose and 1 mM IPTG and plated in top agar. Resistant plaques were picked and propagated on the same strain under the same induction conditions. Candidate recombinant phages were screened by PCR using primers outside the homology arms to confirm *acrVIA1* insertion into the phage genome and to exclude plasmid contamination. An internal PCR targeting the wild-type locus was used to monitor residual WT phage. Serial passages and screening were repeated until no WT amplicon was detectable.

### Transposition infections for pooled TnSeq

Transposition strains and control strains were grown overnight from glycerol stocks in LB supplemented with either 50 μg/mL gentamicin (V1, V2 donors) or 250 μg/mL carbenicillin (V3 donor) at 37 °C with shaking. Overnight cultures were subcultured 1:100 into LB containing 1 mM IPTG (to induce transposase expression), the appropriate antibiotic, and 10 mM Mg^2+^ and grown for 2 h at 37 °C with shaking.

Log-phase cultures were adjusted to OD600 = 0.5 in 3 mL volumes. Bacteria were infected with 10 μL of 2 × 10^4^ PFU/mL ΦKZ at an MOI of approximately 2.22 × 10^-7^ and incubated for 16 h at 37 °C with shaking. The next day, 2 mL of each culture were centrifuged at 16,000 × g for 2 min to pellet unlysed cells. Supernatants were transferred to new tubes, mixed with 100 μL chloroform, and incubated at room temperature with shaking for 10 min. After centrifugation at 10,000 × g for 2 min, supernatants were transferred to fresh tubes, extracted once more with 100 μL chloroform, clarified, and stored at 4 °C.

For the genome-wide pooled TnSeq library, 10 independent transposition reactions done using PAO1 *attTn7:mariner-sfCherry-nlp1 +* p20T-V3 were performed and processed as described above.

### Cas13a counter-selection of transposed phages

Transposition output phages were first titered by serial dilution spot assays on Cas13 non-targeting and dual-targeting strains, as well as in the presence of *jukAB*, to verify identity and exclude contaminants. Typical output titers were ∼4 × 10^10^ PFU/mL.

To enrich for transposed phages, PAO1 *attTn7::cas13aLse* + p30T crRNA^55/120^ strain was grown to OD_600_ ∼1.8 in 3 mL LB and infected at an approximate MOI of 5 using 400 μL of the ∼1 × 10^10^ PFU/mL transposition output. Counter-selection cultures contained 1 mM IPTG, 0.3% arabinose, 50 μg/mL gentamicin, and 10 mM Mg^2+^ and were incubated overnight at 37 °C with shaking. Phages were harvested the following day by chloroform treatment as described above and stored at 4 °C. Serial dilution spot assays and full-plate infections on the counter-selection strain confirmed enrichment and Cas13 resistance, and titers were typically ∼2 × 10^10^ PFU/mL.

For pooled TnSeq, enriched phages from the ten reactions were combined in equal PFU amounts.

### Library preparation for transposon insertion sequencing

#### Phage DNA isolation

Genomic DNA was purified from enriched phage lysates using SDS/proteinase K treatment followed by column purification. Briefly, 50–200 μL of high-titer phage lysate were mixed 1:1 with lysis buffer (final concentration: 10 mM Tris-HCl pH 8.0, 10 mM EDTA, 100 μg/mL proteinase K, 100 μg/mL RNase A, 0.5% SDS), incubated at 37 °C for 30 min and at 55 °C for 30 min, and then processed with the DNA Clean & Concentrator kit (Zymo Research). DNA concentration was measured by spectrophotometry.

#### Illumina library construction and nested PCR

Transposon junction libraries were prepared using the NEBNext Ultra II FS DNA Library Prep Kit for Illumina with modifications to enrich for transposon–genome junctions. To generate indexed i7 adaptors, oAWC190 was pre-annealed separately with oAWC191, 192, 194, or 195 in 10 mM NaCl at 200 μM. For each pair, 12 μL of each primer were mixed, heated to 95 °C for 5 min, and cooled to 12 °C at 0.1 °C/sec. Annealed oligos were diluted to 15 μM and stored at −20 °C.

For fragmentation, 1,000 ng of phage DNA in 26 μL 10 mM Tris-HCl pH 8.0 were mixed with 7 μL Ultra II FS Reaction Buffer and 2 μL Ultra II FS Enzyme Mix, vortexed, and incubated at 37 °C for 24 min to generate 100–250 bp fragments, followed by 65 °C for 30 min and hold at 4 °C. I7 adapters were ligated by adding 30 μL Ultra II Ligation Master Mix, 1 μL Ligation Enhancer, and 2.5 μL of 15 μM pre-annealed adaptor, followed by incubation at 20 °C for 15 min.

For size selection, the reaction was brought to 100 μL with 10 mM Tris-HCl pH 8.0, and 40 μL AMPure XP beads (Beckman Coulter) were added. After 10 min at room temperature and 5 min on a magnetic rack, the supernatant was transferred to a new tube. An additional 20 μL AMPure XP beads were added, incubated 10 min, and the beads washed twice with 200 μL 80% ethanol. Beads were air-dried for 3–5 min and eluted in 16 μL water. Fifteen microliters of eluate containing size-selected DNA were carried forward.

To amplify transposon junctions, 15 μL of size-selected DNA were mixed with 25 μL NEBNext Ultra II Q5 Master Mix, 5 μL 10 μM oAWC196, and 5 μL 10 μM oAWC224 (total 50 μL). PCR was performed with: 98 °C for 30 s; 12 cycles of 98 °C for 10 s, 65 °C for 75 s; 65°C for 5 min; hold at 12 °C. PCR products were cleaned with AMPure XP beads (45 μL beads per 50 μL reaction), washed twice with 80% ethanol, and eluted in 16 μL water (15 μL carried forward).

A second PCR to further enrich junctions and attach i5 adaptors was performed using 15 μL eluate and 35 μL of master mix (25 μL Q5 Master Mix, 5 μL 10 μM oAWC199, and 5 μL of 10 μM oAWC200, 201, 202, or 203). Products were cleaned with AMPure XP beads as above, eluted in 33 μL water, and 30 μL were retained.

Library quality was assessed using the Qubit 1× dsDNA HS Assay (expected 20–50 ng/μL) and Agilent D1000 ScreenTape (expected size ∼300–350 bp).

### Illumina sequencing

Libraries were sequenced on an Illumina MiSeq i100 instrument using the MiSeq i100 Series 25M Reagent Kit (100-cycle; 20126567). Indexed libraries were pooled and diluted with water to ∼4 nM (Qubit-based). A 10x loading concentration of 0.8 nM in 100 μL Resuspension Buffer was prepared according to the manufacturer’s instructions.

A separate indexed sample for ΦKZ whole-genome sequencing was prepared at 0.8 nM and included as a diversity spike-in. The loading mix consisted of 75 μL pooled library and 25 μL spike-in. Runs were configured with 8 cycles for index 1, 8 cycles for index 2, 100 cycles for read 1, and 0 cycles for read 2. Quality control and demultiplexing were performed in Illumina BaseSpace, and FASTQ files for each index were exported for downstream analysis.

### Insertion sequencing analysis

Demultiplexed FASTQ reads were filtered for those containing the transposon inverted repeat sequence marking the insertion junction (GACCGGGGACTTATCAGCCAACCTGTTA). Retained reads were trimmed to keep the last 10 nt of the inverted repeat (CAACCTGTTA) plus 35 nt of downstream genomic sequence.

Trimmed reads were mapped to the reverse complement of the ΦKZ reference genome (NC_004629) using the TraDIS toolkit (bacteria_tradis). Insertion sites supported by ≤2 reads were removed before essentiality analysis. Gene-level insertion statistics were obtained using tradis_gene_insert_sites on both raw and filtered insertion datasets; for filtered data, insertions within the first and last 10% of each annotated ORF were excluded. Fitness-conferring genes were annotated using the tradis_essentiality.R script. TraDIS outputs were further processed and visualized in R.

### Plaque assays

Unless otherwise noted, plaque assays were performed at 37 °C. Strains were grown overnight in LB at 37 °C (EcoRI experiments were typically ∼13 h). For spot assays, 100 μL of overnight culture were mixed with 3–5 mL of 0.4% LB top agar supplemented with Mg^2+^ and poured onto LB + Mg^2+^ plates. After solidification, ΦKZ or derivative phages were serially diluted and 2 μL of each dilution were spotted on the lawn. Plates were incubated overnight at 37 °C.

For complementation assays with the ΦKZ *nlp1::acrVIA1* phage, 150 μL of overnight culture were mixed with 10 μL of phage at the desired dilution and 3.5 mL of 0.35% top agar with Mg^2+^, poured on LB + Mg^2+^ plates containing 50 μg/mL gentamicin, and incubated overnight at 30 °C.

For cold-sensitivity assays, plates were incubated for 48 h at 18 °C. A water-filled plate was placed on top of assay plates to provide weight and limit evaporation.

### qPCR for CBASS experiments

Strains were grown overnight at 37 °C and diluted 1:100 into fresh LB the next day. Cultures were grown for 6 h, and ΦKZ was added at MOI ∼0.5. At each time point, 100 μL of culture were harvested, washed once with 1× PBS, and pellets were snap-frozen on dry ice and stored at −20 °C.

Genomic DNA was extracted using the Zymo Genomic DNA kit. Approximately 1 ng of DNA was used as input for qPCR with primer pairs 15F/15R (phage) and rpoD_F/rpoD_R (bacterial chromosomal control). Phage DNA levels were first normalized to bacterial DNA (ΔCt), and relative phage DNA abundance was then calculated relative to t = 0.

### Adsorption and replication assays

Overnight cultures were diluted 1:100 into fresh LB and grown at 37 °C for ∼2.5 h to OD₆₀₀ ≈ 0.6. ΦKZ (10^4^ PFU) was added to each culture. For adsorption measurements, 100 μL of culture were sampled at t = 15 min, mixed with chloroform to lyse cells, and titrated to determine unadsorbed phage. For replication assays, 100 μL of culture were collected at t = 2 h, lysed with chloroform, and titrated on PAO1 to quantify total phage yield.

### Competition assays

Mutant phages carrying *acrVIA1* were first verified to form plaques on both a non-targeting strain (PAO1 *attTn7::cas13aLse* + p30T^empty^) and a Cas13 single-guide strain PAO1 *attTn7::cas13aLse* + p30T crRNA^120^, while wild-type ΦKZ formed plaques only on PAO1 *attTn7::cas13aLse* + p30T^empty^. Full-plate infections of mutant and wild-type phage on PAO1 *attTn7::cas13aLse* + p30T^empty^ were used to generate starting stocks.

Overnight cultures of PAO1 *attTn7::cas13aLse* with and without guide RNA and were grown at 37 °C. For competition, PAO1 *attTn7::cas13aLse* + p30T^empty^ was subcultured 1:100 and grown to OD₆₀₀ ≈ 0.6. Mutant and wild-type phage were mixed at an approximate 1:1 ratio and added to PAO1 *attTn7::cas13aLse* + p30T^empty^ at a total MOI of 1 in 150 μL volumes in 96-well plates. Infection was monitored over 4 h in a plate reader.

The initial ratio of mutant to wild-type phage was determined at t = 0 by full-plate infection of PAO1 *attTn7::cas13aLse* + p30T^empty^ (total phage) and PAO1 *attTn7::cas13aLse* + p30T crRNA^120^(mutant-only) at appropriate dilutions for plaque counting. After 4 h, chloroform was added to each well and incubated for ∼15 min, and lysates were clarified by centrifugation. The final ratio of mutant to wild-type phage was determined by titration on PAO1 *attTn7::cas13aLse* + p30T^empty^ and PAO1 *attTn7::cas13aLse* + p30T crRNA^120^.

The **Competitive Index (CI)** was calculated as:

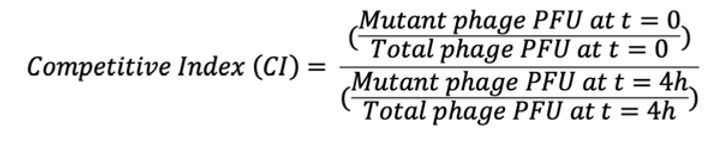

A CI of 1 indicates no change in mutant frequency during the assay (mutant fitness equivalent to wild type), CI < 1 indicates a competitive disadvantage, and CI > 1 indicates a competitive advantage.

## QUANTIFICATION AND STATISTICAL ANALYSIS

For TnSeq analysis, insertions supported by ≤2 reads were excluded and the first and last 10% of each ORF were masked prior to essentiality calling (results in Table S2). Essential and fitness-conferring genes were identified using the TraDIS essentiality pipeline (tradis_essentiality.R). For calculation of total insertions in *orf*s, all reads were considered (results in Table S3). Graphical analysis related to TnSeq were all performed in R.

Plaque assays, adsorption/replication assays, qPCR experiments, and competition assays were typically performed with at least three biological replicates unless otherwise indicated in figure legends. Statistical analysis (e.g., calculation of mean, standard deviation, and, where appropriate, significance testing) was performed in GraphPad Prism; details of tests are indicated in the corresponding figure legends.

### Key resources table

| REAGENT or RESOURCE | SOURCE | IDENTIFIER |
| --- | --- | --- |
| **Antibodies** |  |  |
| **Bacterial and virus strains** |  |  |
| PAO1 p30T<sup>empty</sup> | Qiu *et al.*, 2008<sup>3</sup> |  |
| PAO1 p30T<sup>244-FLAG</sup> | Fossati, Mozumdar *et al.*, 2023<sup>4</sup> |  |
| PAO1 *attTn7::empty* | This paper |  |
| PAO1 *attTn7::jukAB* | Yuping *et al.*, 2025<sup>5</sup> |  |
| PAO1 *attTn7::CBASS* | Huiting *et al.*, 2023<sup>6</sup> |  |
| PAO1 *attTn7::CBASS* + p30T<sup>empty</sup> | Huiting *et al.*, 2023 |  |
| PAO1 *attTn7::CBASS* + p30T<sup>nlp1</sup> | This paper |  |
| PAO1 *attTn7::CBASS* + p30T<sup>acb2</sup> | Huiting *et al.*, 2023 |  |
| PAO1 *attTn7::empty* + p30T<sup>empty</sup> | This paper |  |
| PAO1 *attTn7::empty* + p30T<sup>nlp1</sup> | Huiting *et al.*, 2023 |  |
| PAO1 *attTn7::empty* + p30T<sup>acb2</sup> | Huiting *et al.*, 2023 |  |
| PAO1 p30T<sup>Gb135</sup> | This paper |  |
| PAO1 p30T<sup>Gb136</sup> | This paper |  |
| PAO1 p30T<sup>Gb144</sup> | This paper |  |
| PAO1 *attTn7::mariner-sfCherry-nlp1* | This paper |  |
| PAO1 *attTn7::mariner-sfCherry-nlp1* + p30T-V1 | This paper |  |
| PAO1 + p30T-V1 | This paper |  |
| PAO1 *attTn7::mariner-sfCherry-nlp1* + p30T-V2 | This paper |  |
| PAO1 + p30T-V2 | This paper |  |
| PAO1 *attTn7::mariner-FLAG* + p20T-V3 | This paper |  |
| PAO1 *attTn7::mariner-sfCherry-nlp1* + p20T-V3 | This paper |  |
| PAO1 + p20T-V3 | This paper |  |
| PAO1 *attTn7::cas13aLse* + p30T<sup>nlp1</sup> crRNA<sup>55/120</sup> | This paper |  |
| PAO1 *attTn7::cas13aLse* + p30T<sup>empty</sup> | Guan *et al.*, 2022<sup>2</sup> |  |
| PAO1 *attTn7::cas13aLse* + p30T crRNA<sup>120</sup> | Guan *et al.*, 2022 |  |
| PAO1 *attTn7::cas13aLse* + p30T crRNA<sup>55</sup> | Guan *et al.*, 2022 |  |
| PAO1 *attTn7::cas13aLse* + p30T crRNA<sup>55/120</sup> | This paper |  |
| PAO1 *attTn7::cas13aLse* + p30T crRNA<sup>86start</sup> | This paper |  |
| PAO1 *attTn7::cas13aLse* + p30T crRNA<sup>95cleav</sup> | This paper |  |
| PAO1 *attTn7::cas13aLse* + p30T crRNA<sup>97start</sup> | This paper |  |
| PAO1 *attTn7::cas13aLse* + p30T crRNA<sup>97cleav</sup> | This paper |  |
| PAO1 *attTn7::cas13aLse* + p30T crRNA<sup>99start</sup> | This paper |  |
| PAO1 *attTn7::cas13aLse* + p30T crRNA<sup>119start</sup> | This paper |  |
| PAO1 *attTn7::cas13aLse* + p30T crRNA<sup>162dual</sup> | This paper |  |
| PAO1 *attTn7::cas13aLse* + p30T crRNA<sup>163dual</sup> | This paper |  |
| PAO1 *attTn7::cas13aLse* + p30T crRNA<sup>177start</sup> | This paper |  |
| PAO1 *attTn7::cas13aLse* + p30T crRNA<sup>303start</sup> | This paper |  |
| Env201 | This paper |  |
| PA1032 | This paper |  |
| Phages |  |  |
| ΦKZ | Alan Davidson Lab | AF399011 |
| ΦKZ *orf298::Tn* | This paper |  |
| ΦKZ *orf302::Tn* | This paper |  |
| ΦKZ_*orf152::Tn* | This paper |  |
| ΦKZ *nlp1::acr* | This paper |  |
| ΦKZ *orf2::acr* | This paper |  |
| ΦKZ *orf95::acr* | This paper |  |
| ΦKZ *orf93::acr* | Guan *et al.*, 2022 |  |
| ΦKZ *orf105::acr* | This paper |  |
| ΦKZ *orf244::acr* | This paper |  |
| ΦKZ*orf184::acr* | Li, Xiao, Federova...<br>*et al.*, 2024<sup>7</sup> |  |
| ΦKZ*orf241::acr* | Yuping *et al.*, 2025 |  |
| JBD67 | Alan Davidson Lab | NC_042135.1 |
| JBD67Δ*acb2* | Huiting *et al.*, 2023 |  |
| **Biological samples** |  |  |
| **Chemicals, peptides, and recombinant proteins** |  |  |
| Gentamicin | RPI |  |
| Carbenicillin | BPBio |  |
| IPTG | Fisher |  |
| Chloroform | Alta Aesar |  |
| Proteinase K | Thermofisher Scientific |  |
| RNaseA | Omega |  |
| **Critical commercial assays** |  |  |
| Luna qPCR master mix | New England Biolabs | M3003X |
| NEBNext Ultra II FS DNA Library Prep Kit | New England Biolabs | E7805 |
| NEBuilder HiFi DNA Assembly Master mix | New England Biolabs | E2621L |
| DNA Clean and Concentrator Kit | Zymo Research | D4034 |
| AMPure XP Beads | Beckman Coulter | A63880 |
| Qubit 1x dsDNA HS Assay Kit | Thermofisher Scientific | Q33230 |
| Agilent D1000 ScreenTape and Reagents | Agilent Technologies | 5067-5582<br>5067-5583<br>5067-5602<br>5067-5586 |
| **Deposited data** |  |  |
| **Experimental models: Cell lines** |  |  |
| **Experimental models: Organisms/strains** |  |  |
| *Pseudomonas aeruginosa* strain PAO1 |  |  |
| **Oligonucleotides** |  |  |
| **See Table S1 for details** |  |  |
| **Recombinant DNA** |  |  |
| p30T<sup>empty</sup> | Qiu *et al.*, 2008 |  |
| p30T crRNA<sup>120</sup> | Guan *et al.*, 2022 |  |
| p30T crRNA<sup>55</sup> | Guan *et al.*, 2022 |  |
| p30T crRNA<sup>55/120</sup> | This paper |  |
| p30T crRNA<sup>86start</sup> | This paper |  |
| p30T crRNA<sup>95cleav</sup> | This paper |  |
| p30T crRNA<sup>97start</sup> | This paper |  |
| p30T crRNA<sup>97cleav</sup> | This paper |  |
| p30T crRNA<sup>99start</sup> | This paper |  |
| p30T crRNA<sup>119start</sup> | This paper |  |
| p30T crRNA<sup>162dual</sup> | This paper |  |
| p30T crRNA<sup>163dual</sup> | This paper |  |
| p30T crRNA<sup>177start</sup> | This paper |  |
| p30T crRNA<sup>303start</sup> | This paper |  |
| p30T V1 | This paper |  |
| p30T V2 | This paper |  |
| p20T V3 | This paper |  |
| p30T<sup>244-FLAG</sup> | Fossati, Mozumdar *et al.*, 2023 |  |
| p30T<sup>nlp1</sup> | Huiting *et al.*, 2023 |  |
| p30T<sup>acb2</sup> | Huiting *et al.*, 2023 |  |
| p30T<sup>Gb135</sup> | This paper |  |
| p30T<sup>Gb136</sup> | This paper |  |
| p30T<sup>Gb144</sup> | This paper |  |
| *pUC18-mini-Tn7-mariner-sfCherry-nlp1* | This paper |  |
| *pUC18-mini-Tn7-attTn7::mariner-FLAG* | This paper |  |
| pTNS3 | Choi et al., 2007 |  |
| **Software and algorithms** |  |  |
| GraphPad PRISM | GraphPad Software |  |
| RStudio | R Foundation |  |
| TraDIS Toolkit | Barquist *et al.*, 2016<sup>8</sup> |  |
| **Other** |  |  |

## Supplemental Figure Legends

**Figure S1.**
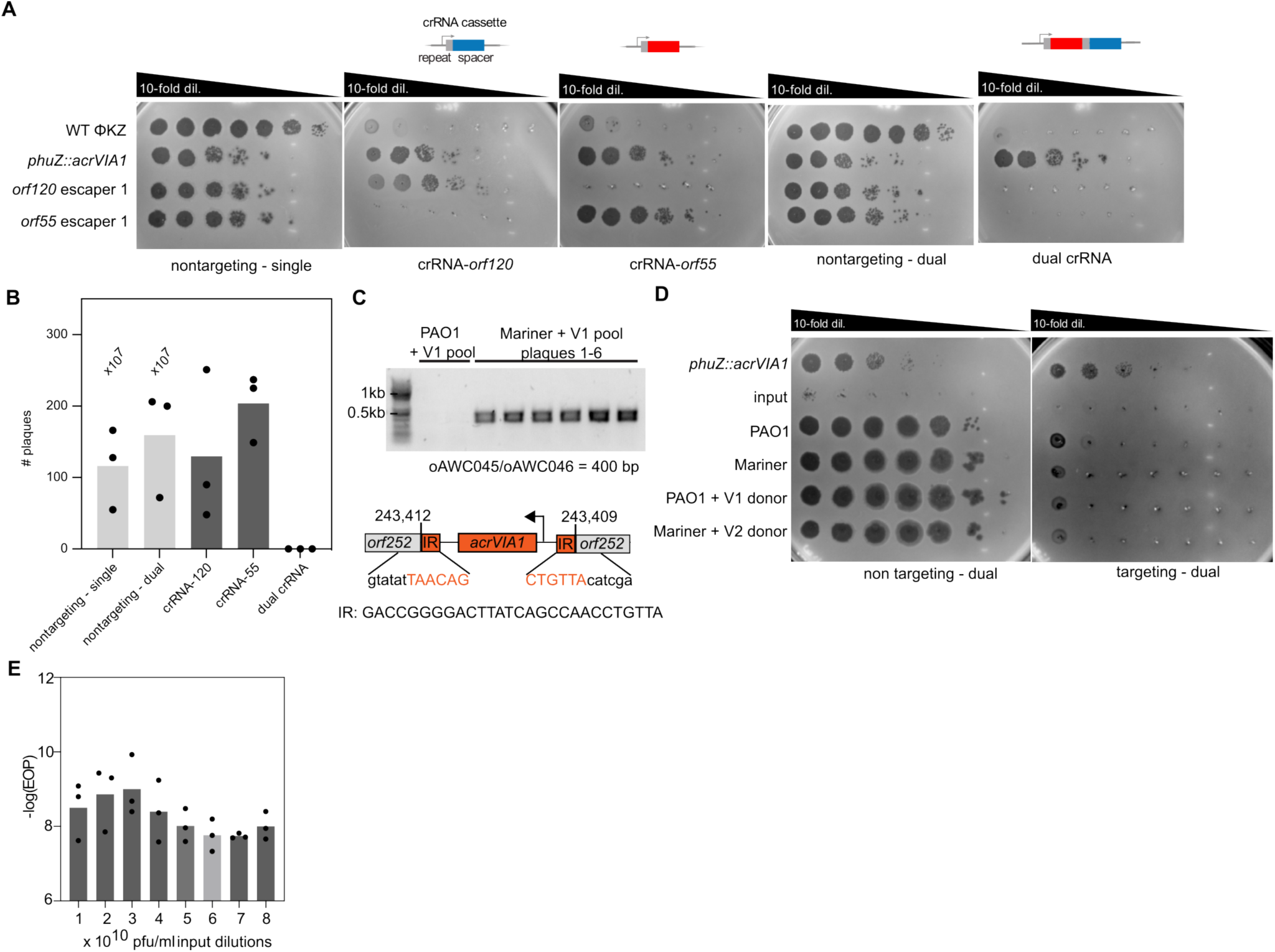
Construction and validation of Cas13 dual-guide counter-selection system. (A) Plaque assays of wild-type ΦKZ, ΦKZ *phuZ*::*acrVIA1*, and single-guide escapers on targeting and non-targeting Cas13 strains. (B) Escape frequencies of single versus dual Cas13a crRNAs targeting ΦKZ. (C) Agarose gel PCR verification of *acrVIA1* insertions in transposed plaques. Low-transposition control pools are shown for comparison. (D) Plaque assays of transposition reactions across targeting and non-targeting hosts. (E) Quantification of transposition efficiency as a function of input ΦKZ titer (pfu/mL).

**Figure S2.**
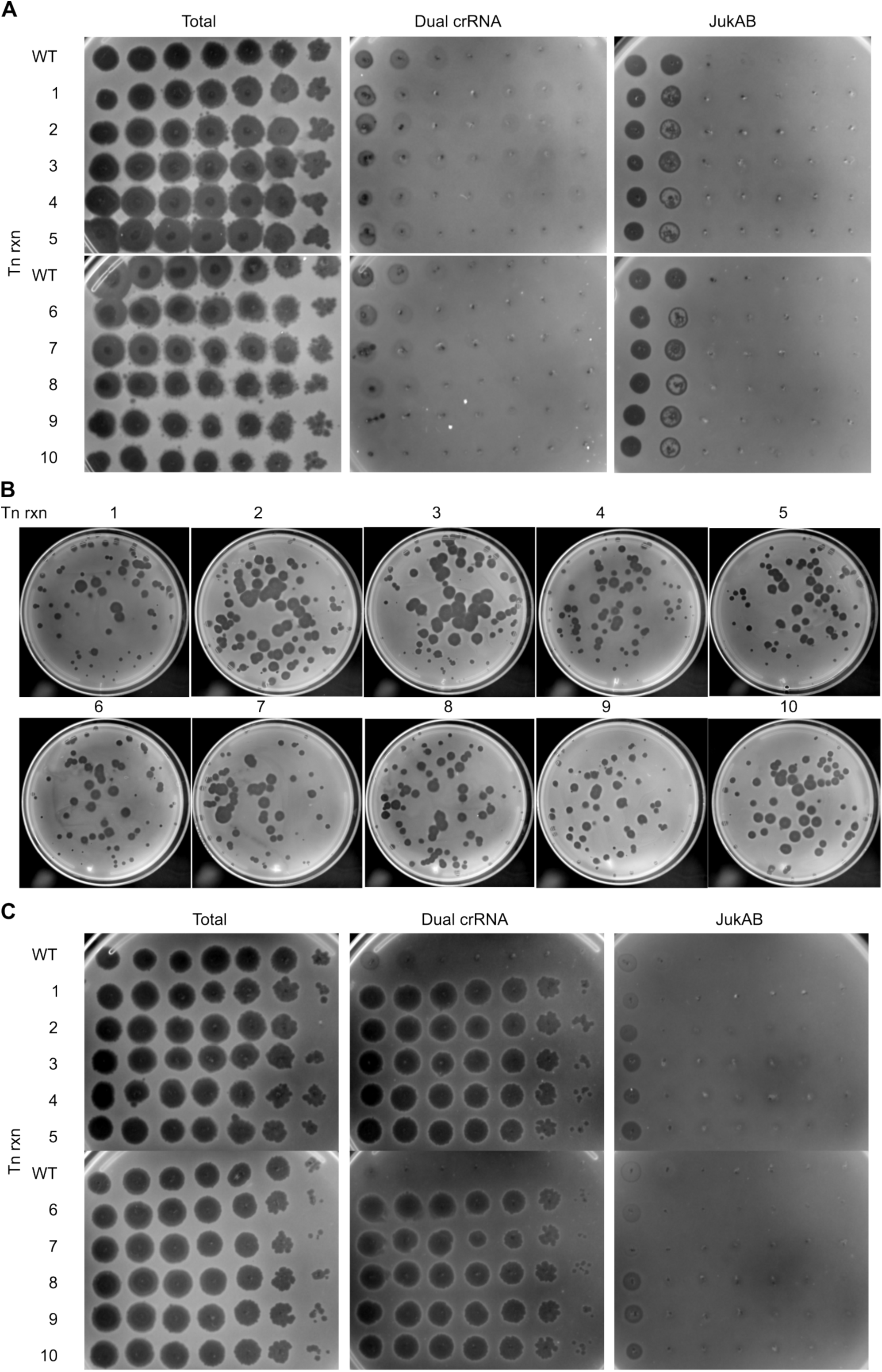
Characterization of individual transposition reactions before and after enrichment. (A) Plaque assays comparing ten independent transposition reactions across selection conditions: no selection, dual Cas13a targeting, and JukAB. Wild-type ΦKZ is shown as control. (B) Full-plate infections (20 μL input) of each transposition reaction. (C) Post-enrichment plaque assays confirming selective recovery of transposed ΦKZ under Cas13a counter-selection.

**Figure S3.**
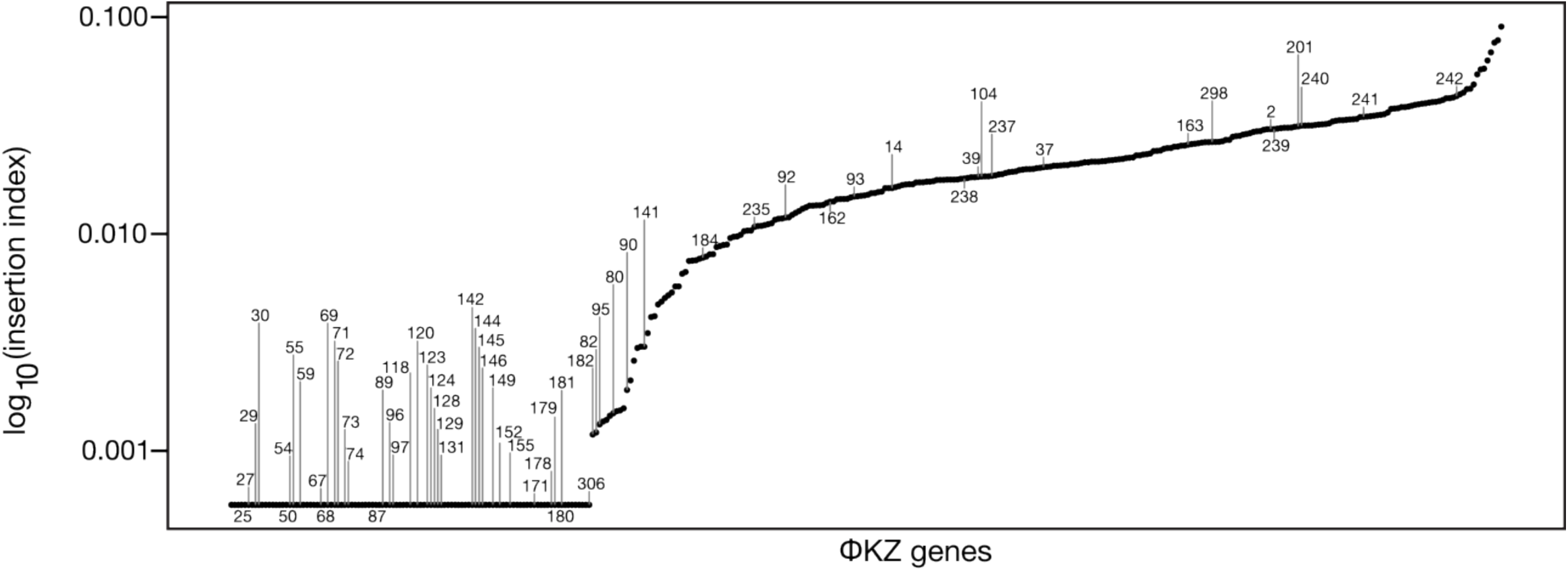
Rank-ordered distribution of ΦKZ gene fitness scores. Cumulative ranking of ΦKZ genes based on transposon insertion frequency and essentiality scoring.

**Figure S4.**
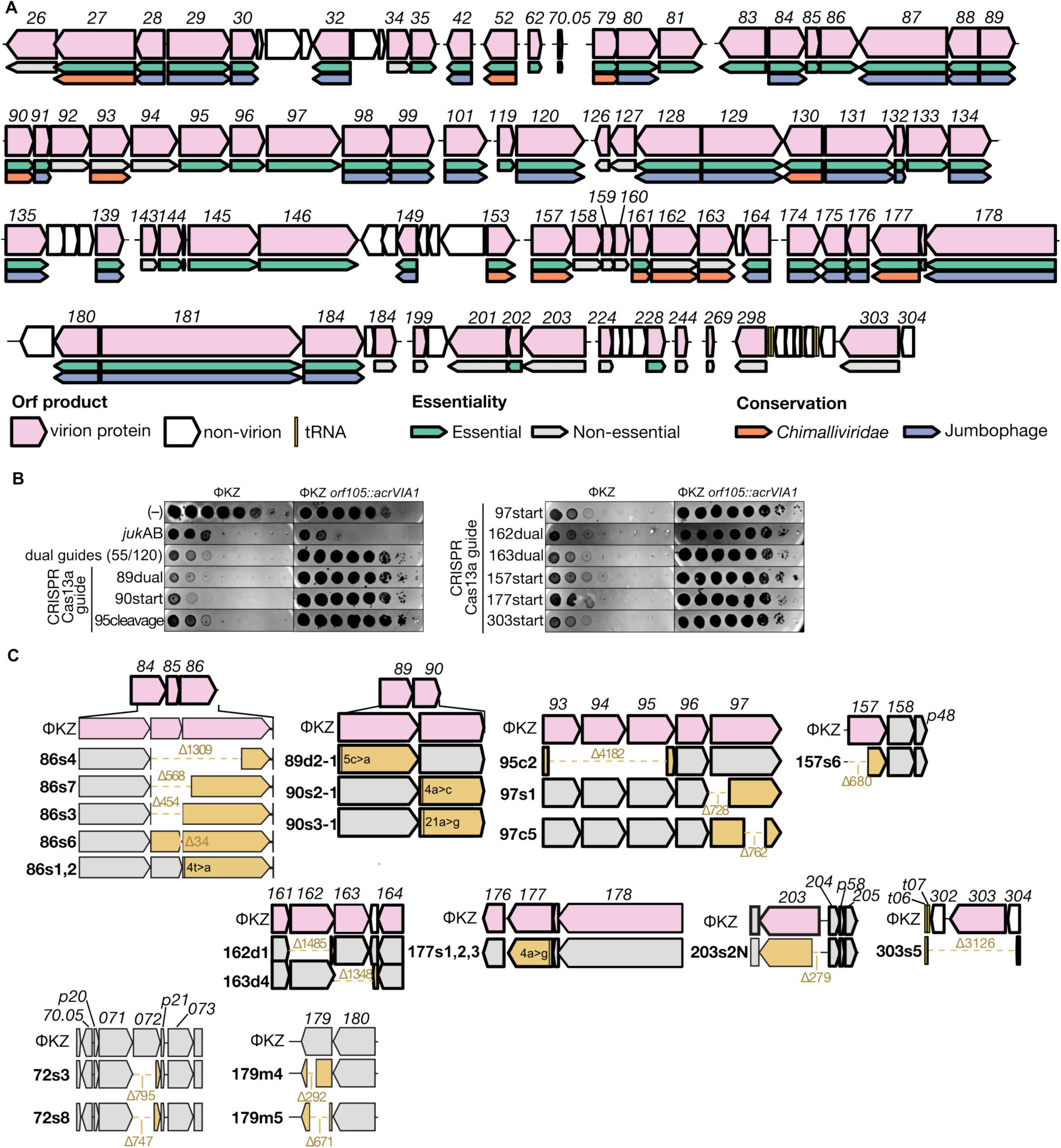
Cas13 escaper phages acquire large genomic deletions. (A) Schematic overview of genes targeted by Cas13. Protein function, essentiality, and conservation across jumbo phages and *Chimalliviridae* are indicated. (B) Plaque assays showing targeting of wild-type ΦKZ and ΦKZ *orf105*::*acrVIA1* (*acrVIA1* control) with crRNAs directed genes encoding injected structural proteins. The Cas13a dual guide construct targeting *orf55* and *120* are included as control. (C) Schematic representation of deletion events identified in Cas13 escaper genomes. The annotations start (s), dual (d) and cleavage (c) refer to the position of guide targeting. Numbers shown indicate the base number within the *orf* that had mutations. Top: virion, bottom: non-virion.

**Figure S5.**
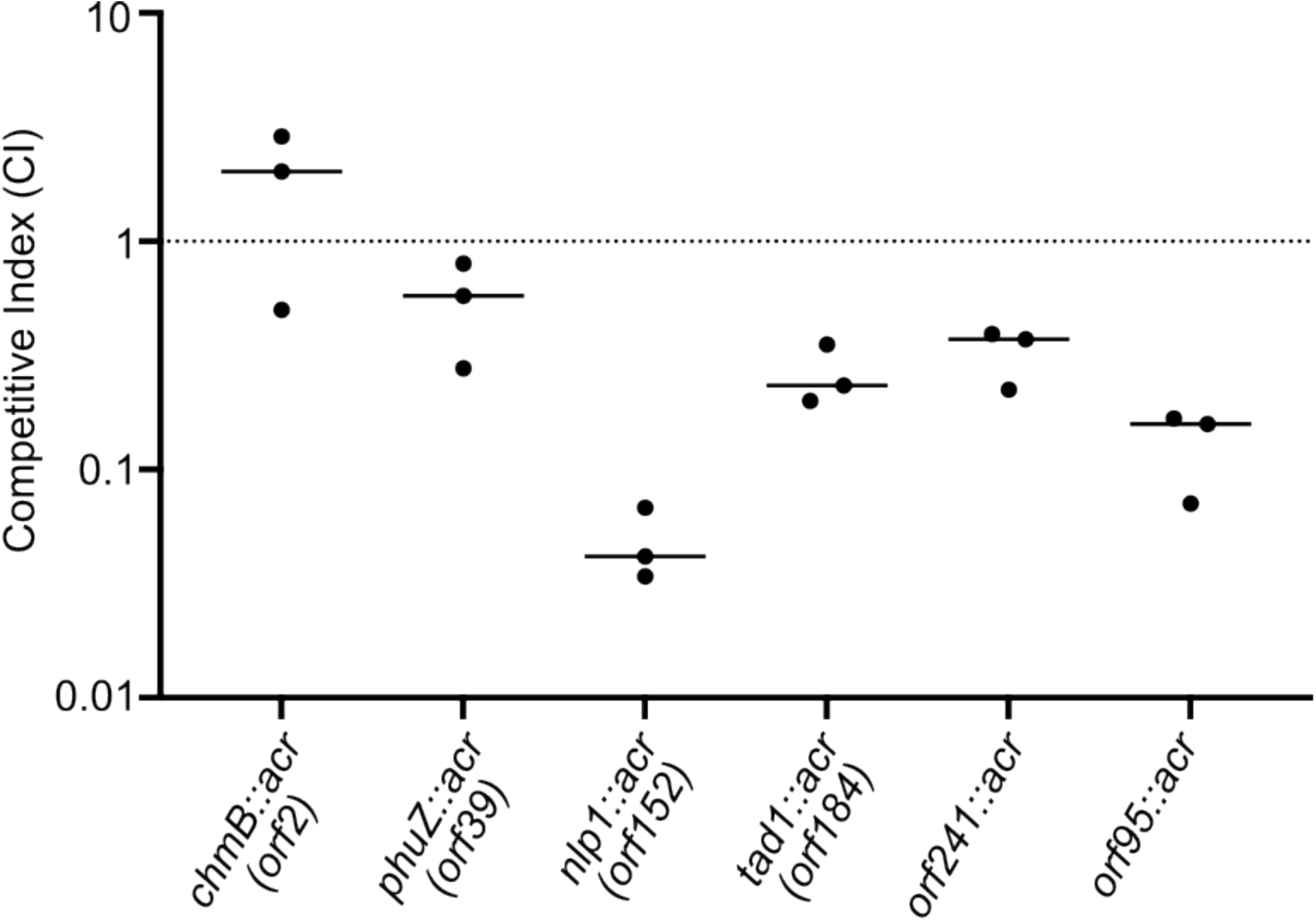
Competitive index of ΦKZ mutants. Competitive fitness of ΦKZ mutants carrying *acrVIA1* insertions in different genes, compared with wild-type phage.

**Figure S6.**
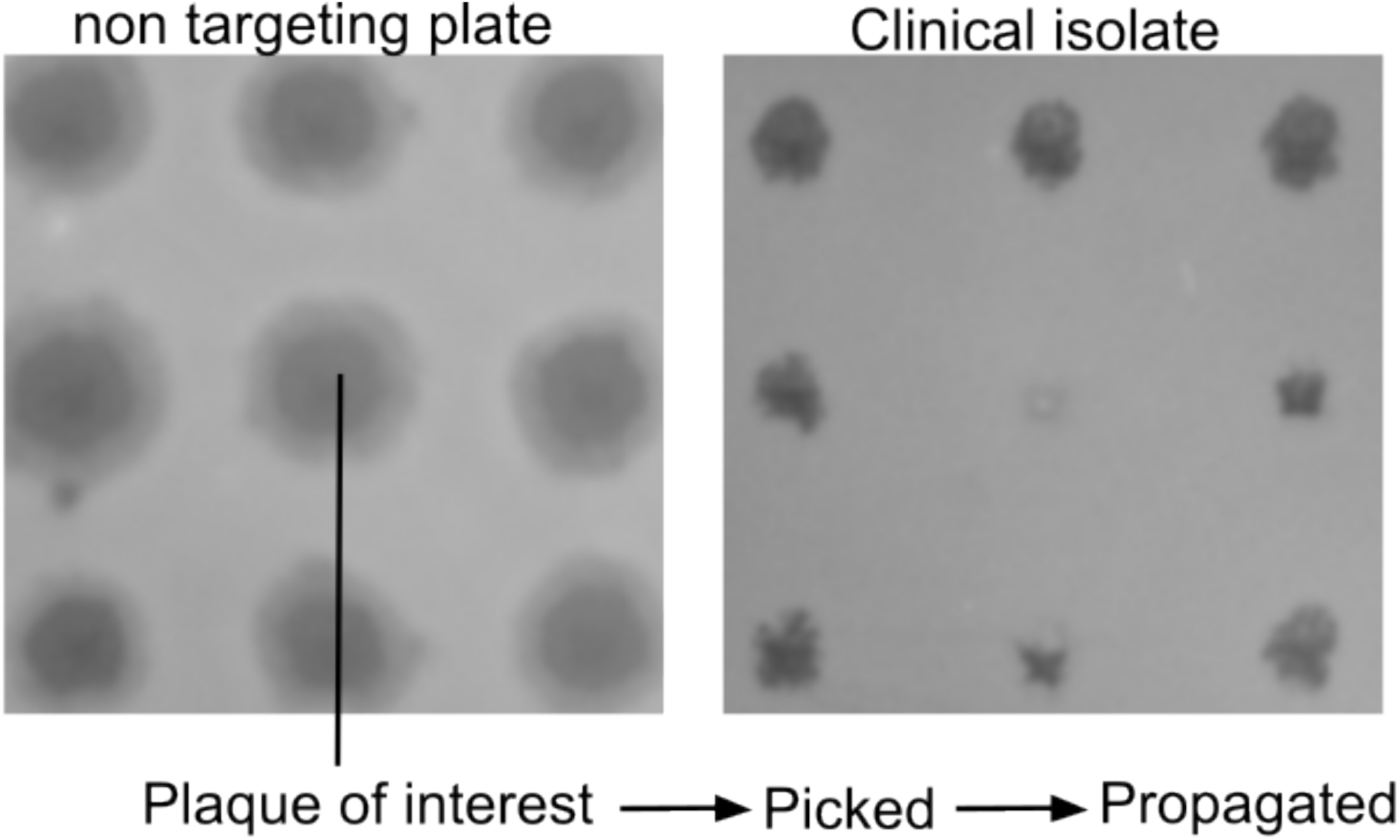
Representative plaques selected for downstream phenotypic follow-up. Example images of plaques from arrayed ΦKZ mutants pinned onto *P. aeruginosa* lawns.

**Figure S7.**
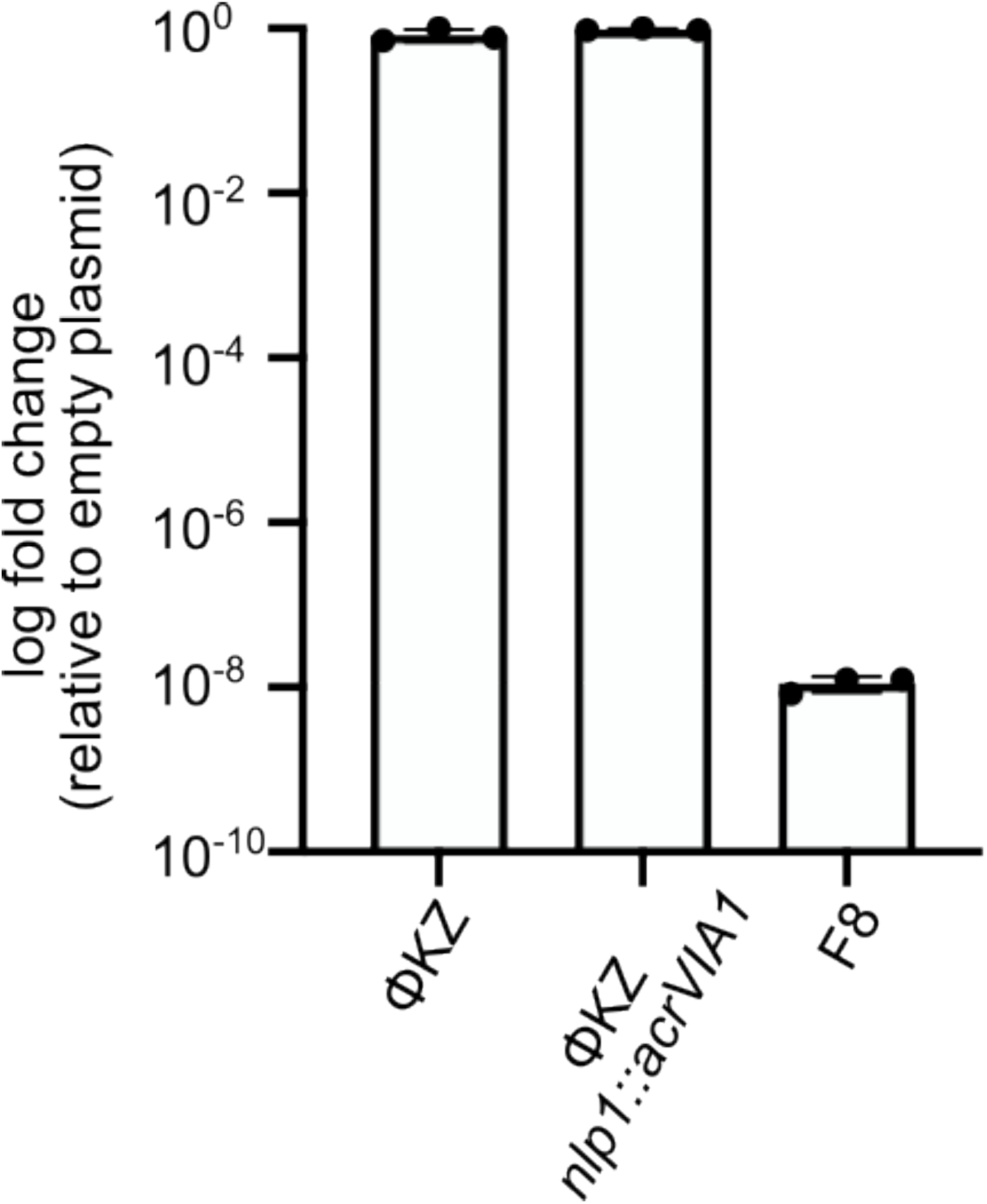
*In vivo* EcoRI targeting of ΦKZ and ΦKZ *nlp1::acrVIA1.* Quantification of *in-vivo* EcoRI restriction of wild-type and ΦKZ *nlp1::acrVIA1*. F8 phage is included as an EcoRI-sensitive control unrelated to ΦKZ.

**Figure S8.**
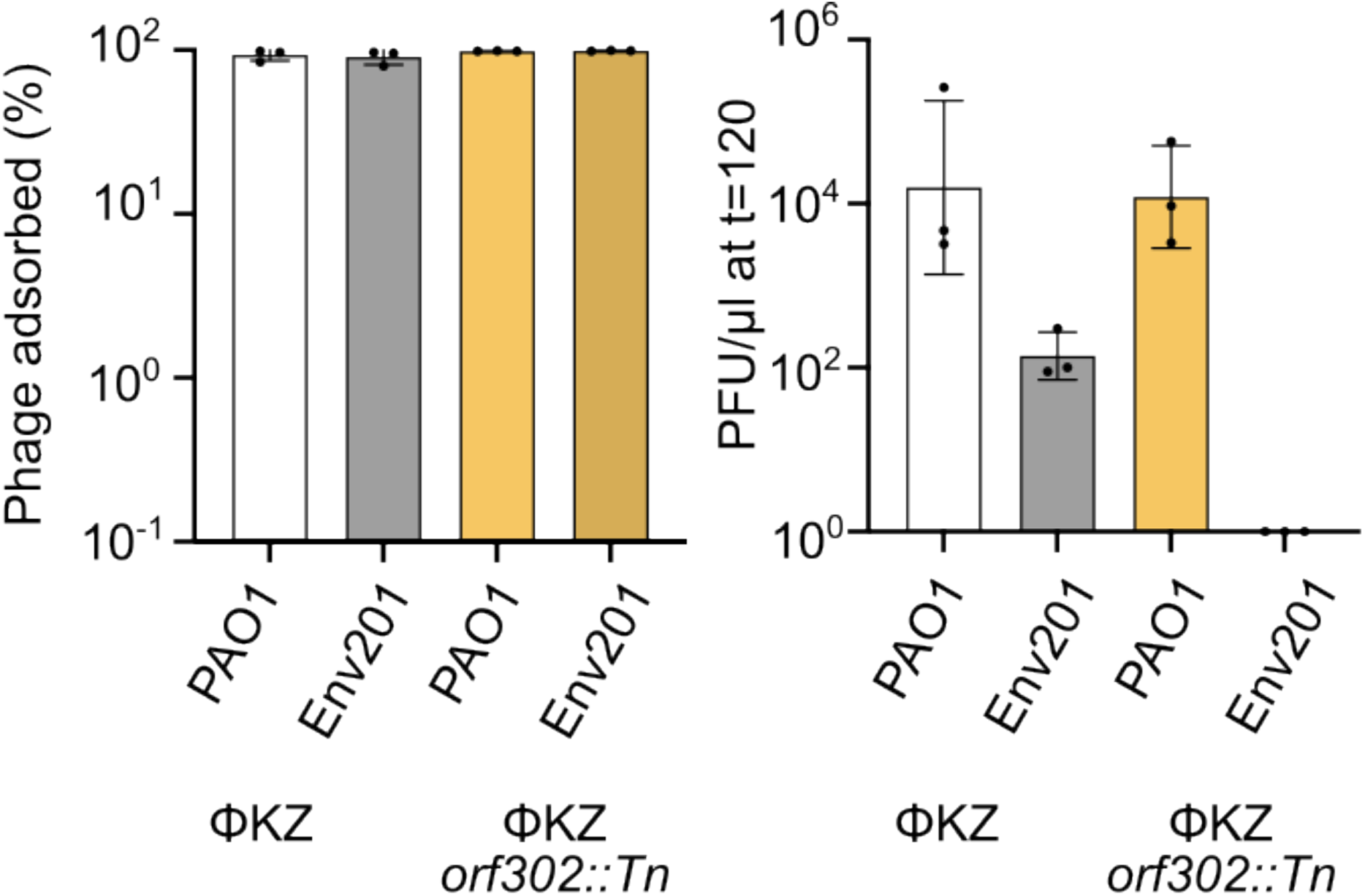
Adsorption and replication kinetics of ΦKZ *orf302::Tn.* Adsorption and 2-hour replication profile of *orf302::Tn* compared to wild-type ΦKZ.

